# Human hepatic stellate cells orchestrate the accumulation and function of CD103^+^ tissue-resident CD8^+^ T-cells in liver fibrosis

**DOI:** 10.64898/2025.11.28.691193

**Authors:** George E. Finney, Daniel Brown Romero, Ginevra Pistocchi, Bethany H. James, Scott P. Davies, Stephanie Kucykowicz, Anandita Mathur, Keshav Rao, Kexin Kong, Jacky Tam, Juliet Luft, Niamah A. Nishan, Giulia Lupo, Walid Al-Akkad, Maria Castanho Martins, Andrew Hall, Brian R. Davidson, Massimo Pinzani, Krista Rombouts, Alberto Quaglia, Mala K. Maini, Prakash Ramachandran, Laura J. Pallett

## Abstract

Tissue-resident memory cells (T_RM_) contribute to protective and pathogenic responses in the liver, yet precisely how hepatic T_RM_ adapt and integrate cues from the underlying stroma and extracellular matrix (ECM) in chronic liver disease (CLD) has yet to be fully defined. Here we describe a role for activated myofibroblast-like hepatic stellate cells (HSCs) in the accumulation and *in situ* localisation of CD8^+^ T_RM_ in the CLD liver. Activated HSCs drive a program of tissue residence in activated, tissue-infiltrating CD8^+^ T-cells in a TGFβ-dependent manner. We show upregulation of CD103, ECM-binding integrins and adhesion molecules driven by TGFβ which together contribute to the sequestration of T_RM_ within the ECM-rich fibrotic niche. *Ex vivo*, hepatic CD103^+^ T_RM_ correlate with the extent of ECM deposited, express an altered repertoire of co-stimulatory and co-inhibitory receptors, transcriptional regulators of cellular exhaustion and produce less proinflammatory mediators upon TCR engagement in CLD than in health. Through expression of several co-inhibitory ligands, we further demonstrate the potential for activated HSCs to acquire an immunomodulatory phenotype and limit the capacity of CD103^+^ T_RM_ to produce anti-viral and anti-tumour mediators upon antigen encounter. Finally, we demonstrate that strategies to block such regulatory pathways, including the PD1:PD-L1/PD-L2 axis, have the potential to restore the antigen-specific effector function of tissue-compartmentalised CD103^+^ T_RM_ and thus contribute to improving the effectiveness of local immunosurveillance in CLD.

**One Sentence Summary:** Activated hepatic stellate cells characteristic of liver fibrosis orchestrate an accumulation of a CD103^+^ T_RM_ population with a reduced capacity for antigen-specific effector function in human CLD.

## INTRODUCTION

The liver is home to a population of tissue-resident CD8^+^ T-cells (CD8^+^ T_RM_) that contribute to dichotomous localised responses^1,2^. On the one hand, they provide highly efficient anti-viral, anti-tumour protective immunity^3–5^ and on the other they have the potential for tissue destruction and immunopathogenesis^6–8^. Through expression of integrins and adhesion molecules, hepatic T_RM_ reside in liver sinusoids, extending pseudopodia-like dendrites to probe the underlying parenchyma for perturbations^2,9–11^. Like those in many other tissues, upon detection of a pathogen or malignancy, hepatic CD8^+^ T_RM_ rapidly secrete proinflammatory cytokines such as IFNγ and TNF and other effector molecules stored in cytotoxic granules to eliminate, or control, infected or transformed cells^3,4,12–14^. However, despite a core transcriptional signature shared by anatomically distinct T_RM,_ recent studies have demonstrated how T_RM_ undergo specific phenotypic and functional adaptations in response to a range of environmental cues once in a tissue niche^15–17^. Beyond environmental cues a CD8^+^ T_RM_ phenotype and function can be further shaped by interactions and exchanges with other cells^12,18^.

An emerging consensus has placed stromal cells at the centre of many pathological processes ranging from inflammation and fibrosis to altered immunity and cancer^19^. Like CD8^+^ T_RM_, stromal cells undergo functional specialisation within a niche. Stromal cells can subsequently exhibit either a matrix-producing, contractile phenotype promoting tissue remodelling, or an immunomodulatory phenotype capable of shaping T cell function, migration and retention. Such regulation of local T-cells is in part through chemokine gradients and ECM-dependent sequestration, but also through expression of inhibitory receptors and the production of modulatory cytokines and chemokines. Stromal cells can for example upregulate PD-L1 to reduce CD8^+^ T cell-mediated immunopathology and promote immune tolerance^20^ and secrete soluble mediators that recruit immunoregulatory population such as Tregs and myeloid-derived suppressor cells that subsequently limit antigen-specific T-cells^21,22^.

Injured HSCs, as a result of excessive consumption of dietary components or alcohol, differentiate from vitamin-A storing quiescent cells into proliferative, fibrogenic myofibroblast-like cells producing high levels of extracellular matrix and TGFβ^23,24^. Given TGFβ’s established role as a regulator of T_RM_ biology^25^, the ability of stromal cells to shape T-cell responses and the functional dichotomy in T_RM_ responses depending on the context^2^, we here explored how hepatic T-cells are shaped by HSCs using CLD as an *in vivo* ‘model system’ whereby myofibroblast-like HSCs significantly alter the tissue niche.

We present a comprehensive phenotypic, functional and transcriptional analysis of hepatic CD8^+^ T_RM_, revealing an accumulation of CD103^+^ T_RM_ in CLD, orchestrated by activated HSCs in an ECM- and TGFβ-dependent manner. In CLD CD103^+^ T_RM_ exhibit a diminished potential for antigen-specific effector function in part due to the expression of immunomodulatory ligands on interacting HSCs. Blockade of regulatory pathways such as PD-L1/PD-L2 can re-establish effective immunosurveillance in CLD by restoring responses with the potential for frontline protection against pathogens and malignancy.

## RESULTS

### Hepatic CD103^+^ T_RM_ increase in abundance in human CLD, trapped within the fibrotic niche

To understand the impact of CLD on CD8^+^ T_RM_ in the human liver, we isolated intrahepatic leukocytes (IHL) from resected or explanted tissues from 137 individuals (Fig.1a). Flow cytometric analysis of hepatic CD8^+^ T-cells using the prototypic residency markers CD69 and CD103 (Fig.1b; gating in <u>Supp.Fig.1a</u>) revealed an increased frequency of CD69, CD103 co-expressing T_RM_ in the CLD liver compared to non-CLD, control tissues - when analysed as a proportion of CD8^+^ T-cells (Fig.1b-c) or total IHL (<u>Supp.Fig.1b</u>). In contrast, the frequency of CD69^+^CD8^+^ T-cells lacking CD103, known to contain long-lived resident cells^26^, significantly decreased in the CLD liver (Fig.1b). When broken down by CLD aetiology, hepatic CD103^+^ T_RM_ were highly abundant in the livers of individuals with metabolic-dysfunction associated steatotic liver disease (MASLD) and alcohol-related liver disease (ALD; Fig.1d). Likewise, image-based quantitative immunofluorescence microscopy enumerating CD103^+^CD8^+^ T-cells *in situ* in an independent cohort (Fig.1e) confirmed an increased percentage of CD8^+^ T-cells co-expressing CD103, and a corresponding increase in the absolute number of CD103^+^CD8^+^ T-cells in CLD (Fig.1f).

**Figure 1:**
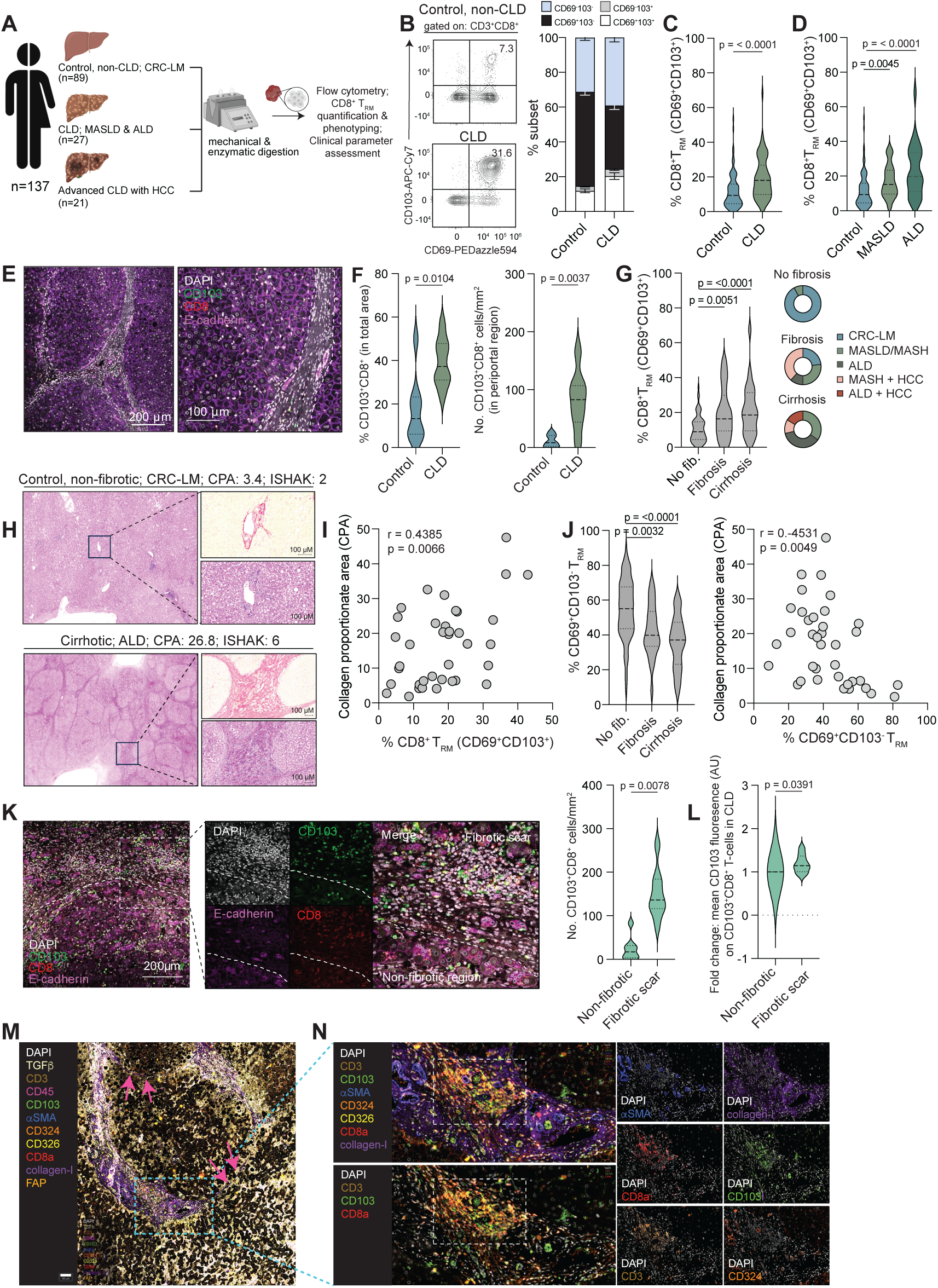
*CD103^+^ T_RM_ increase in abundance in the fibrotic liver.* **a)** Experimental schematic depicting the cohort of tissues used for flow cytometric assessment of the hepatic CD8^+^ T-cell compartment. **b)** Representative flow cytometry plots and summary data showing CD69/CD103 co-expression: defining liver infiltrating, CD69^-^CD103^-^ non-T_RM_ (pale blue), CD69^+^CD103^-^ (black) and CD69^+^CD103^+^ (white) CD8^+^ T_RM_ in healthy, non-CLD livers (Control; samples without clinical indication, or histological evidence of liver disease [n=89]) and CLD livers (n=56). **c)** Frequency of hepatic CD69^+^CD103^+^ T_RM_ identified in health and CLD and **d)** after further categorisation of the CLD cohort by aetiology: n=22 metabolic dysfunction-associated steatotic liver disease (MASLD) and n=26 alcohol related liver disease (ALD). **e)** Representative immunofluorescence staining for CD103 (green), CD8 (red), DAPI (white) and E-cadherin (purple). Right panel showing magnified view. **f**) Percentage CD8^+^ T-cells expressing CD103 observed in the total area analysed and number of CD103^+^CD8^+^ T-cells/mm^2^ in the periportal region in non-CLD (Controls; n=8) and CLD (n=7). **g**) Frequency of CD69^+^CD103^+^ T_RM_ identified by flow cytometry categorised by independent histological assessment of fibrosis stage: no evidence of fibrosis (Healthy), mild-moderate fibrosis (Fibrosis) or advanced fibrosis (Cirrhosis). Pie charts depict aetiology distribution per category. **h)** Representative picrosirius red (PSR) and haematoxylin and eosin (H&E) stains for a non-fibrotic, control and cirrhotic, ALD liver. **i**) Correlation of the frequency of hepatic CD103^+^ T_RM_ with collagen proportionate area (CPA; n=37). **j)** Frequency of CD69^+^CD103^-^ T_RM_ categorised by histological assessment of fibrosis severity as in **g** and correlation with CPA. **k)** Representative immunofluorescence imaging as in **e** of a cirrhotic ALD tissue section and the corresponding summary data delineating number and distribution of CD103^+^CD8^+^ T-cells in the non-fibrotic parenchyma compared to the fibrotic scar. White dotted lines separate the region of scarring from the non-fibrotic region. **l**) Fold change in CD103 staining intensity on CD103^+^CD8^+^ T-cells in the fibrotic scar compared to the non-fibrotic region. **m**) Representative images of a fibrotic tissue stained with an 11-plex MACSima multiplex panel. Blue box identifying the fibrotic scar. **n**) Magnified view and single stain images of the ECM-rich fibrotic scar region from **m** depicting CD3^+^CD8^+^CD103^+^ T_RM_ within a cellular infiltrate (white box) surrounded by collagen-I. Violin plots show median ± quartiles. Each tissue sample was stained and processed independently.

Combining all tissues, hepatic CD103^+^ T_RM_ accumulation was not a feature of sex or age (<u>Supp.Fig.1c-d</u>), nor did it correlate with the extent of liver inflammation when assessing serum transaminases (alanine transaminase [ALT] and aspartate aminotransferase [AST]; <u>Supp.Fig.1e-f</u>). Rather, the frequency of CD103^+^ T_RM_ associated with the underlying fibrotic response in those with CLD. Categorisation by independent histological assessment into three groups: no evidence of fibrosis, mild-moderate fibrosis (fibrosis) or advanced fibrosis (cirrhosis) revealed an increased CD103^+^ T_RM_ frequency with fibrosis stage, regardless of aetiology (Fig.1g). In support of CD103^+^ T_RM_ increasing with fibrosis severity, the frequency of hepatic CD103^+^ T_RM_ by flow cytometry correlated with collagen deposition in the liver as measured by assessment of the collagen proportionate area (CPA) calculated from picrosirius red staining of adjacent samples^27,28^ (Fig.1h-i), and the ISHAK score, a histological grading system assessing inflammation and fibrosis (<u>Supp.Fig.1g</u>). Conversely, the frequency of CD103^-^ T_RM_ decreased with fibrosis severity and negatively correlated with the CPA (Fig.1j).

To gain a more in-depth view of CD103^+^ T_RM_ abundance and localisation in CLD, combined immunofluorescence imaging with histological assessment of consecutive haematoxylin and eosin (H&E) stained sections delineating the fibrotic scar and uninvolved, non-fibrotic regions (<u>Supp.Fig.1h</u>) revealed an enrichment of CD103^+^ T_RM_ within the fibrotic scar (Fig.1k). Scar-associated CD103^+^ T_RM_ exhibited an increased staining intensity for CD103 in tissue sections (Fig.1l) which was supported by an increased overall mean fluorescence intensity of CD103 staining on T_RM_ in CLD compared to non-CLD controls by flow cytometry (<u>Supp.Fig.1i</u>). Further examination of scar-associated CD103^+^ T_RM_ using MACSima imaging detected CD103^+^ T_RM_ in a cellular infiltrate (dashed white box; <u>Fig.1m</u>) within the scar, surrounded by a dense network of fibrillar collagen (Fig.1n) in an ALD liver with bridging fibrosis (pink arrows; Fig.1m). Here, few CD103^+^CD8^+^ T-cells exited into the non-fibrotic parenchyma (Fig.1m-n).

Taken together we demonstrate that hepatic CD103-expressing CD8^+^ T_RM_ expand in CLD irrespective of aetiology and accumulate within the ECM-rich, fibrotic niche.

### Activated hepatic stellate cells promote CD103^+^ T_RM_ accumulation via the production of ECM proteins and TGFβ

Given the correlation of CD103^+^ T_RM_ with CPA, their increased abundance in the scar, and proximity to alpha smooth muscle expressing (αSMA^hi^) myofibroblast-like cells *in vivo* (Fig.2a) we next examined the potential for HSCs to shape the T_RM_ compartment. Upon injury HSCs transdifferentiate from quiescent vitamin A storing cells into activated αSMA^hi^ myofibroblast-like cells secreting excessive amounts of ECM, matrix metalloproteinases, tissue inhibitors of metalloproteinases and TGFβ^23^. Consequently, we investigated whether TGFβ produced by primary human αSMA^hi^ myofibroblast-like HSCs (<u>Supp.Fig.2a-b</u>) could promote a residency program in activated CD8^+^ T-cells, in part via regulation of *ITGAE* (CD103) expression^25^. To address this, co-cultures of HSCs isolated from non-fibrotic or cirrhotic tissues with TCR- or interleukin-15 (IL-15)-activated PBMC

**Figure 2:**
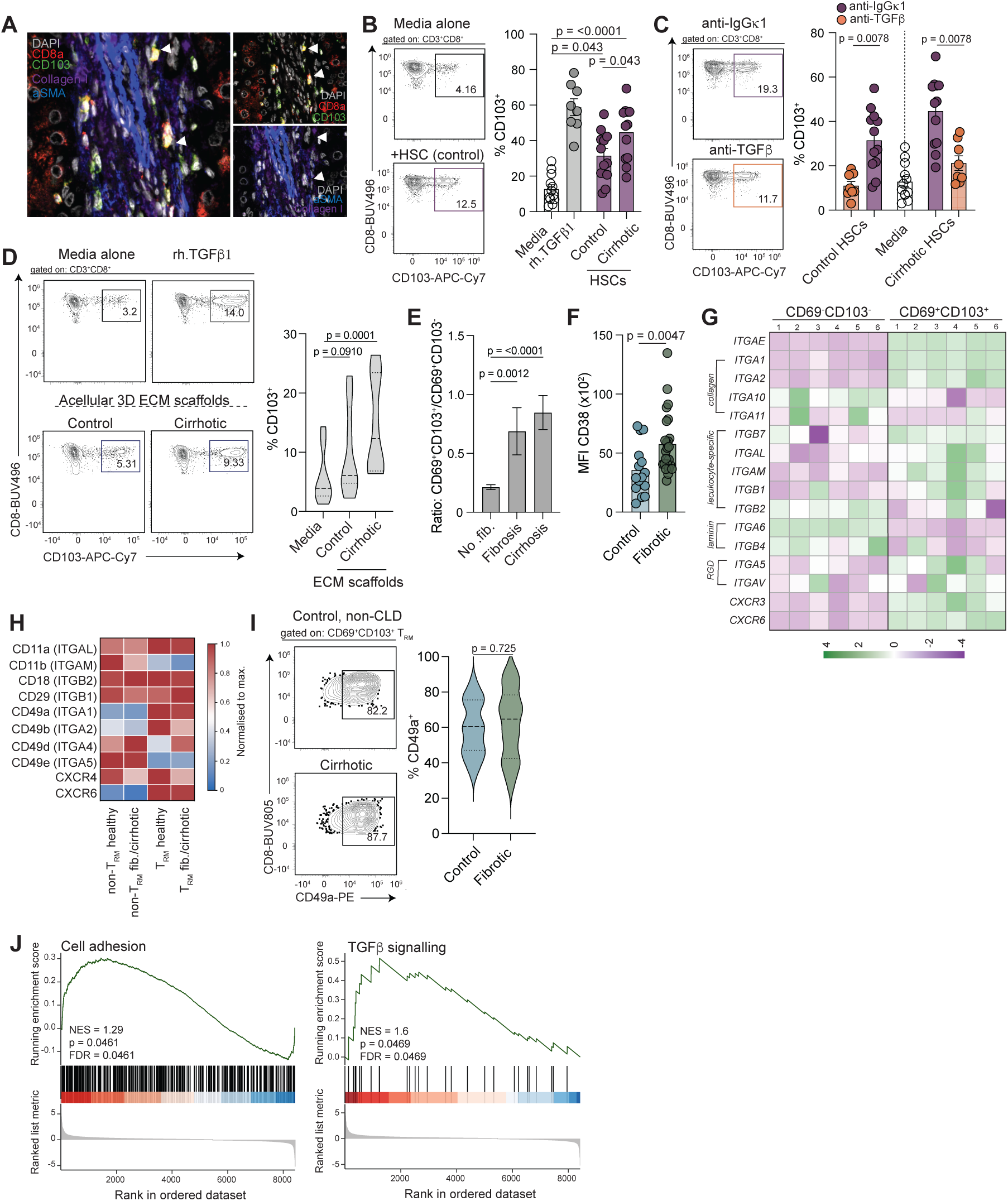
*Activated HSCs orchestrate CD103^+^ T_RM_ accumulation through TGFβ and ECM protein production.* **a)** Representative MACSima images showing CD103 (green), CD8 (red), DAPI (white), collagen-I (purple) and αSMA (blue). **b)** Representative flow cytometry plot and summary data depicting percentage CD103 expression on peripheral TCR-activated CD8^+^ T-cells in media alone (white; n=8-12), after recombinant TGFβ1 exposure (grey; n=8) or co-culture with primary aSMA^hi^ activated myofibroblast-like hepatic stellate cells (HSCs) isolated and cultured from either a non-fibrotic, control or a cirrhotic liver tissue (n=8-12) and **c)** percentage CD103 after co-culture on HSCs in the presence of either an IgG1κ isotype control or pan-TGFβ neutralisation antibody (n=8). **d)** Representative flow cytometry plots and summary data depicting percentage CD103 expression on peripheral IL-15-activated CD8^+^ T-cells in media alone, after recombinant TGFβ1 exposure or co-culture in 3D acellular ECM-scaffolds derived from either a non-fibrotic, control or cirrhotic liver tissue (n=8). **e)** Mean fluorescence intensity (MFI) of CD38 expression on *ex vivo* CD103^+^ T_RM_ from tissues without any clinical indication, or histological evidence of liver disease (Control: blue) and a combined cohort with mild-moderate fibrosis and advanced fibrosis (Fibrotic; green). **f**) Fold change in the frequency of CD103^-^ vs CD103^+^ T_RM_ *ex vivo* in IHL isolated from tissue samples assessed by flow cytometry and categorised by independent histology assessment of fibrosis stage: no evidence of fibrosis (No fib.), mild-moderate fibrosis (Fibrosis) or advanced fibrosis (Cirrhosis). **g)** Bulk RNA sequencing of FACS-sorted *in vitro*-derived IL-15/TGFβ1 CD69^-^CD103^-^ and CD69^+^CD103^+^ T_RM_-like cells (n=6). Heat map depicting ECM-binding integrin and adhesion molecule transcript abundance. **h)** Heat map summarising corresponding protein expression *ex vivo* on CD69^-^CD103^-^ liver-infiltrating, non-T_RM_ and CD103^+^ T_RM_ – data presented are the mean expression of each marker per subset assessed by flow cytometry. **i)** Representative flow cytometry plots and summary data depicting expression of CD49a (ITGA1) on CD103^+^ T_RM_ in tissue without any clinical indication, or histological evidence of liver disease (Control: blue) compared to a combined cohort with mild-moderate fibrosis and advanced fibrosis (Fibrotic; green). **j**) GSEA for genes set relating to TGFβ signalling and cell adhesion. NES, normalised enrichment score; FDR, false discovery rate. Each dot represents an individual PBMC/IHL sample processed independently. Error bars show mean ± S.E.M. Violin plots show median ± quartiles.

(naturally devoid of CD103^3^) were established. Interaction of activated CD8^+^ T-cells with HSCs resulted in the marked upregulation of CD103 (Fig.2b TCR-activated**_**&**_**<u>Supp.Fig.2c</u> IL-15-activated), whereas the addition of a pan-TGFβ neutralising antibody abrogated HSC-mediated CD103 upregulation (Fig.2c). Activated HSCs from cirrhotic tissues further upregulated CD103 compared to those from non-fibrotic control tissues (Fig.2b).

Through association with latent-TGFβ binding proteins, HSC-secreted TGFβ can bind ECM proteins such as fibrillin and fibronectin anchored for proteolytic cleavage^29^. Consequently, we tested whether TGFβ retained within the ECM contributes to the accumulation of CD103^+^ T_RM_ in CLD. For this we used decellularised human liver ECM-scaffolds, an acellular 3D model previously demonstrated to retain native ECM and physiologically relevant concentrations of TGFβ^30–32^. IL-15-activated CD8^+^ T-cells re-populating ECM-scaffolds significantly upregulated CD103 (Fig.2d). In line with the accumulation of CD103^+^ T_RM_ in CLD (Fig.1b-c), the percentage of CD8^+^ T cells expressing CD103 peaked in ECM-scaffolds derived from cirrhotic CLD (Fig.2d). In support of enhanced TGFβ sensitivity in CLD, we observed an increased proportion of CD103^+^ compared to CD103^-^ T_RM_ (Fig.2e), with CD103^+^ T_RM_ expressing more CD38 - an ectonucleotidase previously shown to impact TGFβ responsiveness^33^ - *ex vivo* when isolated from fibrotic tissues (Fig.2f).

We then proceeded to investigate expression of pertinent adhesion molecules and ECM-binding integrins on CD103^+^ T_RM_ to address the postulate that HSC-derived TGFβ contributes to both an accumulation and sequestration in CLD within the ECM-rich scar. Bulk RNA-sequencing of FACS-sorted IL-15/TGFβ1-induced CD69^-^CD103^-^ CD8^+^ non-T_RM_-like and CD103^+^ T_RM_-like cells revealed increased expression of collagen-binding integrins (*ITGA1* and *ITGA2*), RGD-binding integrins that bind an Arg-Gly-Asp tripeptide motif in ECM proteins (*ITGA5* and *ITGAV*) and leukocyte-specific receptors (*ITGAL* and *ITGAM*) on CD8^+^ T-cells expressing CD103 as a result of *in vitro* TGFβ1 exposure (Fig.2g). Of note, TGFβ1 exposure significantly decreased transcript expression of the laminin receptors *ITGA6* and *ITGB4* on CD103^+^ T_RM_-like cells compared to non-T_RM_-like cells. To validate transcriptome observations, we assessed protein expression of several integrin and adhesion transcripts on *ex vivo* CD8^+^ T-cells from IHLs from control, non-fibrotic and fibrotic tissues. In agreement with the transcript, protein expression of CD49a (ITGA1), CD49b (ITGA2) and CD11a (ITGAL) were increased on CD103^+^ T_RM_ compared to non-T_RM_ counterparts (Fig.2h). Whereas protein expression of CD49d (ITGA4) was significantly lower on CD103^+^ T_RM_ than non-T_RM_. Furthermore CD103^+^ T_RM_ protein level expression of CD11b (ITGAM) and CD49e (ITGA5) differed from the transcript in CD103^+^ T_RM_-like cells induced by TGFβ1. Despite ligand availability considerably changing in CLD, protein expression of many ECM-binding integrins remained equivalent on CD103^+^ T_RM_ irrespective of fibrosis severity (Fig.2h-i<u>_&_Supp.Fig.2d</u>).

Finally, to corroborate these findings, we probed a published single cell transcriptomic atlas of the human liver. For this 107,488 non-proliferating CD8^+^ T-cells were re-clustered into eight CD8^+^ T-cell subpopulations (<u>Supp.Fig.2e</u>), identifying a distinct T_RM_ cluster with high level expression of tissue residency genes *ITGAE*, *CXCR6*, *CD69*, *ITGA1, ZNF683* and *RUNX2* (<u>Supp.Fig.2f-g</u>). Differential gene-expression analysis comparing T_RM_ in health and CLD found a total of 1409 differentially expressed genes. Of these genes 1110 were more highly expressed in CLD T_RM_ and 299 more highly expressed in their healthy control counterparts. Gene set enrichment analysis revealed that T_RM_ in CLD showed a specific enrichment of genes associated with enhanced TGFβ signalling and cell adhesion (Fig.2j).

Taken together, these data suggest that exposure to high concentrations of TGFβ produced by HSCs in the fibrotic niche upregulates CD103 on activated, liver infiltrating CD8^+^ T-cells and can elicit the co-expression of ECM-binding integrins that contribute to the sequestration of CD103^+^ T_RM_ in CLD.

### CD103^+^ T_RM_ exhibit an altered co-inhibitory/co-stimulatory receptor profile and reduced potential for antigen-specific responses in CLD

To address the impact of fibrosis on CD103^+^ T_RM_ function we probed the potential for dichotomous protective anti-viral, anti-tumour responses and the capacity to contribute to immune-mediated tissue destruction and pathogenesis^2^. We examined CD103^+^ T_RM_ function testing both TCR-dependent and TCR-independent stimulation of IHL from non-CLD (Control) and a combined cohort of patients with CLD with confirmed fibrosis or cirrhosis (Fibrotic). Validating previous studies^3,12,14,34^, CD103^+^ T_RM_ from the non-fibrotic liver rapidly produced more IFNγ and TNF upon TCR-stimulation compared to liver-infiltrating CD69^-^CD103^-^ T-cells (non-T_RM_; Fig.3a). However, when TCR-stimulated CD103^+^ T_RM_ from the fibrotic liver produced less cytokines than T_RM_ from control livers, instead secreting similar amounts of IFNγ and TNF to their non-T_RM_ counterparts (Fig.3a). Intriguingly, CD103^+^ T_RM_ in fibrosis maintained their remarkable ability to produce high levels of IL-2 (<u>Fig.3a</u>). The functional impairment in T_RM_ in CLD was T_RM-_specific, with non-T_RM_ producing equivalent amounts of cytokines in response to TCR engagement regardless of underlying fibrosis (Fig.3a). Additionally, rather than favouring cytokine production over cytotoxicity like T_RM_ from the non-fibrotic liver (<u>Fig.3a-b</u>), CD103^+^ T_RM_ in fibrosis appeared to more rapidly mobilise cytotoxic granules when TCR-stimulated, and in fact exhibited enhanced cytotoxicity than non-T_RM_ (Fig.3b).

**Figure 3:**
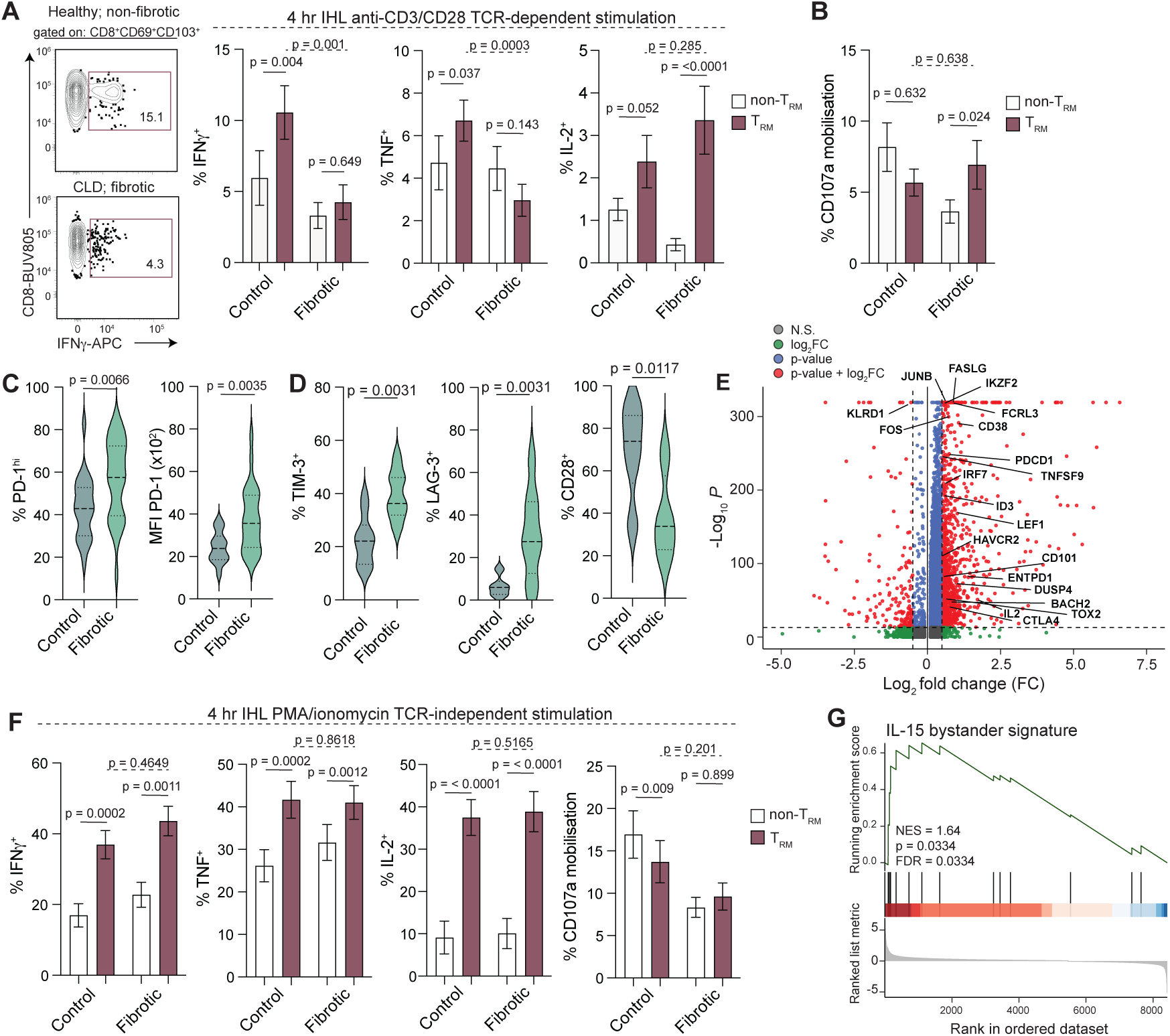
CD103^+^ T_RM_ from the fibrotic liver exhibit a dysfunctional, ‘exhausted’ profile associated with reduced effector function to TCR engagement. Representative flow cytometry plots (IFNγ only) and summary data depicting percentage of CD69^-^CD103^-^ liver-infiltrating, non-T_RM_ (white) and CD103^+^ T_RM_ (red) producing **a)** IFNγ, TNF, IL-2 and **b)** mobilising CD107a^+^ cytotoxic granules after short-term stimulation with anti-CD3/CD28 (n=22 healthy and n=29 fibrotic/cirrhotic) categorised by histological fibrosis stage assigned by an independent histopathologist. **c)** Percentage PD-1^+^ CD103^+^ T_RM_ and density of PD-1 expression on PD-1^+^CD103^+^ T_RM_ and **d**) percentage TIM3^+^, LAG3^+^ and CD28^+^ CD103^+^ T_RM_ identified in tissue samples without any clinical indication, or histological evidence of liver disease (Control) compared to a combined cohort of samples with mild-moderate fibrosis and advanced fibrosis (Fibrotic). **e**) Volcano plot depicting differentially expressed transcripts between T_RM_ (cluster defined in Supp.Fig.2e) in health and CLD. Grey represents genes not differentially expressed (N.S.), green those with a log2 fold change (FC), blue genes with a significant *p* value and red those significantly expressed with a log2 FC. **f**) Percentage CD69^-^CD103^-^ liver-infiltrating, non-T_RM_ and CD103^+^ T_RM_ producing IFNγ, TNF, IL-2 and mobilising CD107a^+^ cytotoxic granules after short-term stimulation with PMA/ionomycin (n=17 healthy and n=21 fibrotic/cirrhotic) categorised as in **a**. **g**) GSEA for core gene relating to IL-15 bystander activation. NES, normalised enrichment score; FDR, false discovery rate. Violin plots show median ± quartiles. Error bars represent mean ± S.E.M.

In line with a reduced potential for antigen-specific effector function, CD103^+^ T_RM_ in fibrosis, expressed an altered co-stimulatory/co-inhibitory receptor profile. Furthering our previous work showing high levels of PD-1 on T_RM_^26,35^, the percentage of PD-1^+^CD103^+^ T_RM_ *ex vivo* significantly increased in the fibrotic liver, with the density of PD-1 on PD-1^+^CD103^+^ T_RM_ significantly elevated (Fig.3c_&_<u>Supp.Fig.3a</u>). Moreover, CD103^+^ T_RM_ from fibrotic livers expressed higher levels of other co-inhibitory receptors including TIM-3 and LAG-3, and markedly reduced levels of the co-stimulatory molecule CD28 (<u>Fig.3d</u>**_**&_<u>Supp.Fig.3b</u>). Corroborating the enhanced expression of individual co-inhibitory receptors, multidimensional analysis revealed signatures likely driving the reduced capacity for TCR-dependent effector function within the pool of CD103^+^ T_RM_. Such signatures included a CD28^lo^TIM-3^hi^ and a PD-1^hi^LAG-3^hi^TIGIT^hi^ signature, which both increased in abundance in fibrosis when confirmed by traditional flow cytometric gating (<u>Supp.Fig.3c-d</u>). Similarly, differential gene expression analysis (false discover rate (FDR) = 0.05 | log (fold change) ≥0.05) of the transcriptional profile of healthy and CLD T_RM_ corroborated the upregulation of genes promoting T-cell dysfunction and cellular exhaustion in CLD. We observed increased expression of co-inhibitory receptors including *PDCD1, HAVCR2, CTLA4, CD101, TNFRSF9* and *ENTPD1* and transcriptional regulators associated with exhaustion such as *IKZF2, ID3, TOX2* and *DUSP4* (Fig.3e). Notably we also observed increased expression of *BACH2*, *JUNB*, *FOS* and *LEF1* and reduced expression of *KLRG1* that may restrain terminal differentiation, enhance survival and longevity of CD103^+^ T_RM_ and contribute to the sustained ability to produce IL-2 to TCR-stimulation observed in CLD^36,37^.

Instead, when exposed to antigen-independent bystander activation using PMA/ionomycin bypassing T-cell receptor engagement, CD103^+^ T_RM_ from the fibrotic liver retained a capacity for rapid effector function – producing high levels of proinflammatory mediators - with no T_RM-_specific functional impairment (Fig.3f). When considering the potential for bystander cytotoxicity, PMA/ionomycin-stimulated CD103^+^ T_RM_ exhibited a similar profile to their non-T_RM_ counterparts (Fig.3f). Moreover, further evaluation of differentially expressed genes in CLD T_RM’s_ with gene-set enrichment analysis revealed an enrichment for a distinct and robust gene signature indicative of innate-like bystander activation in CLD (Fig.3g).

In summary CD103^+^ T_RM_ in CLD present a phenotypic and functional profile indicative of T-cell dysfunction, with increased expression of inhibitory checkpoint molecules, low co-stimulatory receptor expression, and a reduced ability to produce IFNγ and TNF in a TCR-dependent manner. In contrast, CD103^+^ T_RM_ in CLD functionally diverge from those in health with an enhanced potential for bystander, antigen-independent activation and contribution to immune-mediated damage.

### Activated HSC express an altered immunomodulatory profile and contribute to hepatic CD103^+^ T_RM_ dysfunction in CLD

Comparison of candidate immunomodulatory genes expressed by portal fibroblasts, HSCs and activated HSCs, in health and CLD from the single cell RNA-atlas, revealed increased expression of several immunomodulatory ligands or mediators with a capacity to limit T-cell effector function (Fig.4a). Further flow cytometric profiling of our primary αSMA^hi^ myofibroblast-like HSCs confirmed high level protein expression of co-inhibitory PD-L1, PD-L2 and CD155 (Fig.4b**_**&**_**<u>Supp.Fig.4a</u>), with significantly increased expression of PD-L1 on HSCs from CLD compared to non-fibrotic controls (Fig.4b).

**Figure 4:**
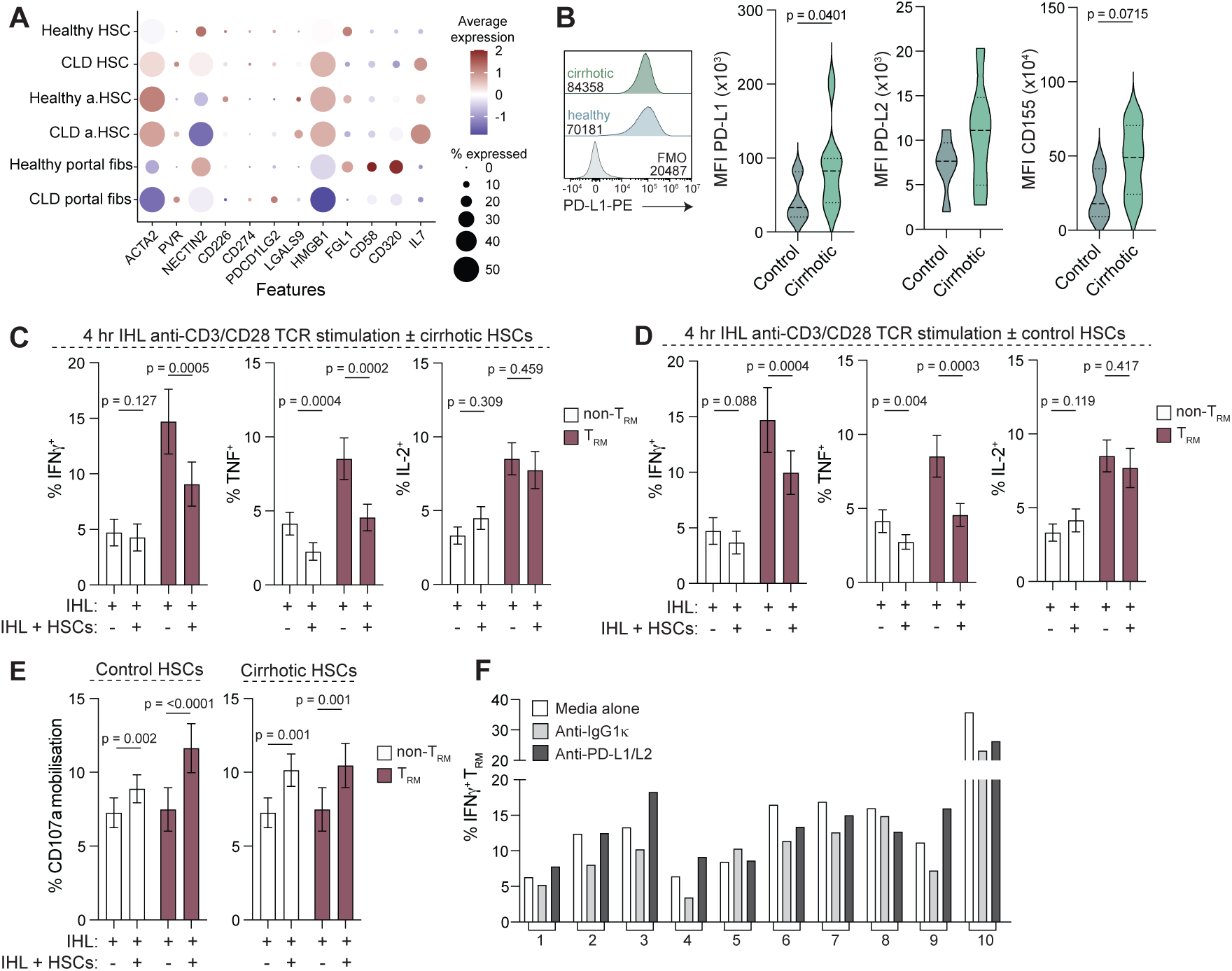
Activated hepatic stellate cells acquire an immunomodulatory phenotype in CLD and contribute to the diminished antigen-specific effector function by CD103^+^ T_RM_. **a**) Dotplot depicting the mean expression of immunomodulatory genes and the percentage of subset expressing them using publicly available RNA-transcriptomic data. **b**) Representative flow cytometry and histogram plots showing surface expression of PD-L1 on HSCs isolated from two donors (one non-fibrotic, control [blue] and one cirrhotic [green]) compared to a fluorescence minus one control (FMO; grey) and summary data showing mean fluorescence intensity (MFI) of surface PD-L1, PD-L2 and CD155 expression on HSCs isolated and cultured from non-fibrotic, control (n=7) and cirrhotic (n=8) donors. Summary data depicting percentage of CD69^-^CD103^-^ liver-infiltrating, non-T_RM_ (white) and CD103^+^ T_RM_ (red) producing IFNγ, TNF or IL-2 after overnight TCR-dependent stimulation with anti-CD3/CD28 (n=11) in the presence or absence of HSC obtained from a **c**) cirrhotic liver or **d**) non-fibrotic, control liver. **e**) Summary data depicting CD69^-^CD103^-^ liver-infiltrating, non-T_RM_ and CD103^+^ T_RM_ CD107a mobilisation in the presence or absence of HSC obtained from the non-fibrotic, control liver (left panel) or cirrhotic liver (right panel). **f**) Summary data depicting percentage of CD103^+^ T_RM_ producing IFNγ after overnight stimulation with anti-CD3/CD28 (n=11) in the presence of HSC pre-incubated with either PD-L1/L2 neutralising antibodies (dark grey) or an isotype control (light grey; IgGκ1) for each individual donor tested. Violin plots showing median ± quartiles. Error bars showing mean ± S.E.M.

As postulated co-cultures of functional CD103^+^ T_RM_ from non-fibrotic tissues with activated HSCs from either non-fibrotic or fibrotic tissues resulted in reduced proinflammatory cytokine production after both a four-hour (Fig.4c-d) and overnight (<u>Supp.Fig.4b</u>) TCR-dependent stimulation. In support of our *ex vivo* finding in CLD, co-cultures of IHLs with HSCs did not impact the percentage of CD103^+^ T_RM_ capable of producing IL-2 (<u>Fig.4c-d</u>). In contrast, upon interaction with HSCs from either tissue source CD103^+^ T_RM_ mobilised CD107a^+^ cytotoxic granules to their plasma membrane without exogenous stimulation, indicative of increasing cytotoxic activity (<u>Fig.4e</u>). Pre-incubation of HSCs with blocking antibodies against an exemplar co-inhibitory pathway - PD-L1 and PD-L2 (<u>Supp.Fig.4c</u>) - prior to the addition of CD103^+^ T_RM_ restored or partially restored the ability of CD103^+^ T_RM_ to produce IFNγ to antigen-dependent stimulation in the majority of individuals tested (Fig.4f). These data suggest a mechanism whereby HSCs characteristic of the cirrhotic liver restrain antigen-dependent cytokine production and instead promote a T_RM_ with enhanced potential for bystander activation.

## DISCUSSION

Here we reveal the impact of pathogenic fibrosis orchestrated by transdifferentiated myofibroblast-like HSCs on hepatic T_RM_. We expose the capacity for HSCs to promote a program of tissue residency in CD8^+^ T-cells patrolling through the liver. As a prominent source of TGFβ, HSCs elicit expression of CD103 (integrin αE) and other ECM-binding integrins and adhesion molecules, including CD49a and LFA-1^9^, that contribute to an accumulation of CD103^+^ T_RM_ in CLD, and their sequestration with the fibrotic niche. Additionally, we report the potential for HSCs to modulate T_RM_ effector function via the expression of a number of immunomodulatory ligands.

Hepatic T_RM_ are not only crucial for homeostasis but essential for protective immunity. Nevertheless, studies in human CHB, and mouse models of MASH and MASH-related HCC called into question their functional role, establishing the potential for T_RM_ ‘autoaggression’ and contribution to immune-mediated damage^6–8^. Therefore, debate has arisen as to whether these apparently opposing roles are mediated by distinct subsets occupying the liver or as a result of differential environmental stimuli^2^. In the skin functionally distinct T_RM_ subsets exist, separated by expression of CD49a^38^, whereas our data suggest altered effector function within a pool phenotypically similar T_RM_. Instead proposing that hepatic T_RM_ integrate and respond, with different effector functionality, to a myriad of additional T-cell extrinsic cues, including contact-dependent interactions with immunomodulatory HSCs, soluble mediators and the underlying ECM.

Corroborating previous studies, we show activated HSCs store latent-TGFβ (mature TGFβ associated to the latent-associated peptide [LAP]) and that the ECM deposited can anchor and make mature TGFβ available to patrolling T-cells. Although not explored, the precise composition of ECM (i.e. the extent of fibrillin and fibronectin deposition^29^) and expression of latent-TGFβ-cleaving integrins such as GARP, avβ8 and avβ6 expressed by resident cells within the sinuosids in CLD likely regulate and control TGFβ avalibility^39–41^ and thus also impact T_RM_ differentiation. Beyond acting as sink for mature TGFβ, the ECM likely also impacts T_RM_ abundance by conveying pro-survival signals via interaction of CD49a with collagen-1 and -IV^26,42^. Interestingly other studies in mice suggest that while T_RM_ in organs like the skin require TGFβ for residency, those in the murine liver (that lack CD103) alternatively use retinoic acid to drive a TGFβ-independent T_RM_-differentiation pathway^15,16^. As HSCs represent a store of vitamin A (retinol) which can be converted into retinoic acid it is tempting to speculate that this may represent a further mediated produced by HSCs capable of shaping hepatic T_RM_. However, in contrast to the mouse, our data suggest that CD8^+^ T_RM_ in the human liver remain highly sensitive to local TGFβ. Supporting this, T_RM_ in the human CLD liver upregulate CD38 – a molecule proposed as a novel regulator of T_RM_ development. Experimental knock-down of CD38 specifically affects the differentiation of CD103^+^ T_RM_ populations impacting sensitivity to TGFβ and expression of TGFβRII^33^.

In addition to their accumulation, our data suggest that activated HSCs in CLD negatively impact the functional capacity of CD103^+^ T_RM_ – limiting responsiveness to antigen-specific stimuli and the production of essential antiviral, anti-tumour mediators. While we propose a direct role for activated myofibroblast-like HSCs in the functional skewing of T_RM_ through the expression of co-inhibitory ligands, it is plausible that these are not the only mechanisms impacting T_RM_ functionality *in vivo*. Although also not investigated here the mobility and parenchymal sampling rate of T_RM_ in regions rich in fibrillar ECM in the CLD liver are likely to be significantly reduced, as reported in the murine uterine tract^43^. Confounding this inability to move freely is the progressive loss of sinusoidal fenestrations during LSEC capillarisation characteristic of fibrosis^11,44^. Such morphological changes raise the possibility that once trapped within the fibrotic niche the ability of CD103^+^ T_RM_ to access and efficiently probe underlying hepatocytes for pathogens or malignancy is substantially hindered, thereby limiting antigen-specific activation. Similarly, it is highly plausible that T-cell expression of mechanosensors enable sensing of environmental stiffness. Exposure to stiff surfaces, akin to those experienced in the fibrotic liver, have the potential to drive cellular exhaustion^45^. With further validation it would be interesting to determine whether the increased co-inhibitory receptor profile observed in individuals with fibrosis represents an adaptation made in response to increasing tissue stiffness. Alternatively, the exhaustion profile exhibited by T_RM_ in CLD may represent an adaptation to balance fibrotic sequelae with protective immunity as seen in the virally infected lung^46^.

Finally, we previously demonstrated the remarkable ability of CD103^+^ T_RM_ to secrete IL-2^3^. Therefore, it is notable that CD103^+^ T_RM_ retain the capacity to produce IL-2 in CLD, when production of other proinflammatory cytokines upon TCR-stimulation is diminished. Thus, raising the possibility that CD103^+^ T_RM_ are somehow protected from full ‘exhaustion’ and upregulate pathways capable of restraining terminal effector differentiation^37^. Taken together with the potential for autocrine IL-2 this may ensure the CLD liver is populated by some, albeit less effective, long-lived memory responses mediating protection upon pathogen re-encounter^37^.

In summary, we demonstrate that highly activated HSCs contribute to the accumulation and functional skewing of human liver T_RM_. The discovery that T_RM_ accumulate irrespective of aetiology but rather as a consequence of fibrogenesis (a process intended to repair and remodel), raises additional questions. For one, exactly how does the specific pattern of ECM laid down impact T_RM_ biology, given the extensive integrin expression profile that actively ‘traps and retains’ T_RM_ within scars? Secondly what is the exact phenotype of the interacting HSCs *in vivo* modulating T_RM_ biology? And thirdly, how will strategies to reverse fibrosis impact the pool of compartmentalised T_RM_, on one hand potentially improving the motility and surveillance of established T_RM_ populations, but on the other if bioavailable mature TGFβ becomes limiting it is possible that novel therapies will inadvertently limit the differentiation of new T_RM_ specificities to emerging threats. Ultimately, we uncover a population in CLD that may be amenable to therapeutic intervention to restore important anti-viral/anti-tumour responses within the liver.

## MATERIALS AND METHODS

### Study samples, ethics and inclusion

Each study participant provided written informed consent before inclusion. Sample storage conformed to the requirements of the *Declaration of Helsinki*, the *Data Protection Act 1998* and the *Human Tissue Act 2004*.

Tissue samples were obtained from either explanted livers obtained after solid-organ transplantation or resected, distal, non-diseased associated margins (NAT) of individuals having surgery for colorectal cancer liver metastases (CRCLM) or hepatocellular carcinoma (HCC; detailed in **Supp.Table1**), surplus to diagnostic requirements. This included individuals with MASLD or MASH and with ALD (individuals whose weekly alcohol consumption exceeded 140g for females or 210g for males). Most liver tissue samples used in this study were obtained through the Tissue Access for Patient Benefit Initiative (TAPB) at The Royal Free Hospital (RFH; approved by the UCL-Royal Free Hospital BioBank Ethical Review Committee references: 11/WA/0077, 16/WA/0289 and 21/WA/0388). Additional FFPE-liver tissue samples were obtained in Birmingham from the Queen Elizabeth Hospital for imaging (approved by the University of Birmingham Ethics Committee reference: 18/WA/0213). These tissues were obtained from explanted livers after solid organ transplantation for ALD, or from healthy donor livers from organs deemed unsuitable for transplantation. In addition, healthy donor blood samples were obtained by local collection (approved by the Royal Free London NHS Foundation Trust and UCL Biobank, Ethical Review Committee reference: 21/WA/0388).

Donors under the age of 18, HIV seropositive donors, HCV RNA positive and donors with high viral titre hepatitis B infection (HBV; DNA >10 IU/mL) were excluded from the study. No further exclusion criterion was used; all available samples with sufficient viable cell yields for the experimental protocol were included. No statistical method was used to predetermine sample size. In addition, no sample randomisation was used in this study; all participants were pseudo-anonymised at time of surgery (before delivery of material to the research team) and sampled *ad hoc*. All tissue samples were subsequently assigned to fibrosis stage/score groups by independent consultant histopathologists (Prof Alberto Quaglia and Andrew Hall; RFH).

### Sample processing & cell isolation

Intrahepatic leukocytes (IHL) were isolated from tissue samples as previously described^47^. Briefly, tissue samples were dissociated and incubated at 37 °C, 5% CO_2_ in HBSS^+/+^ (Life Technologies; Thermo Fisher Scientific) containing 0.002% DNase I (Roche; Sigma-Aldrich; Merck) and 0.02% collagenase IV (Thermo Fisher Scientific). After enzymatic digestion, tissue samples were subjected to mechanical disruption using the GentleMACS (Miltenyi Biotech) dissociator and debris removed by filtration through a 70 µM filter (Griener). Parenchymal cells were removed by centrifugation on a 30% Percoll gradient (GE Healthcare; VWR) followed by leukocyte isolation by a further round of density centrifugation using a Pancoll gradient (PAN Biotech).

Peripheral blood was obtained from healthy controls. PBMC were isolated from heparinised peripheral blood by density centrifugation using a Pancoll gradient.

### Single-cell RNA sequencing analysis

Human single-cell RNA-seq data were obtained from eight published studies^48–55^ and collated into an extended single cell atlas by Papachristoforou, Kong & Colella *et al*. Full details on analysis employed to store and merge data files, the QC filtering procedures and subsampling to obtain the original liver atlas are as previously described. To illustrate the expression patterns of key marker genes across the different lineages, dot plots were generated. These plots provided a clear visualisation of both the average expression levels and the proportion of T-cells expressing each gene within each identified lineage.

The CD8^+^ T-cell object obtained from the original published atlas^56^ was subsetted and saved as an independent S5 object. Subsequent analysis of this object involved FindVariableFeatures, ScaleData, RunPCA, RunUMAP (dims=1:20), FindNeighbors (dims=1:20), FindClusters (resolution=1), and annotation for the DecontXcounts assay. Marker genes were identified using the FindAllMarkers function, applying the same thresholds as those used for the whole atlas. Doublet clusters (expression of multiple lineage markers) and low-quality clusters (median nFeature < 800) were manually labelled and removed iteratively, to ensure that only high-quality cells remained. Final clustering was performed at resolution=0.4 - a manually determined resolution. Annotation of each CD8^+^ T-cell cluster was then performed manually using marker genes lists and literature searches. Upon annotation of the CD8^+^ T_RM_ cluster, the same pipeline was run as above with the exception of the final clustering resolution, which was performed at resolution=0.1.

Differential composition analysis (DCA) was performed on the six selected CD8^+^ T_RM_ clusters using the package *MiloR* (version 2.4). Based on the KNN graph, shared cell neighbourhoods were generated for each lineage using the functions buildGraph and makeNhoods, with varying k, d, and prop values. Differential neighbourhood abundance testing was then conducted using the testNhoods function. For the CD8^+^ T_RM_ subpopulations, cell-type-level comparisons between several groups were conducted: healthy vs. CLD, fibrosis stage (comparing no – minimal fibrosis [F0-1] and advanced fibrosis [F4]) and male vs. female. To ensure robust statistical analysis, we restricted the analysis to that of the two largest CD8^+^ T_RM_ clusters (clusters 0 and 1) containing at least 50 neighbourhoods per cell type.

Statistical comparisons of relative abundances between groups were performed using unpaired two-sample t-tests (Student’s t-test), implemented via the stat_compare_means() function from the ggpubr *R* package. P-values were not adjusted for multiple comparisons. Differential gene expression analysis (DGEA) was performed for CD8^+^ T-cells. FindMarkers (test.use = MAST, only.pos = F, min.pct = 0.01, logfc.threshold = 0.5) was used to perform DGEA between CD8^+^ T_RM_ subclusters 0 and 1. The resulting gene list was further filtered by p_val_adj < 0.01 and avg_log2FC > 0.5.DotPlot, VlnPlot, pheatmap, and EnhancedVolcano were utilised for the visualisation of the results in *R*.

### FACS Sorting and Bulk RNA sequencing of cytokine-induced CD103^+^ T_RM_

PBMC were cytokine induced to derive a T_RM_-like phenotype as previously described^3^. Briefly, 1 x 10^6^ cells/well were incubated with 50 ng/mL IL-15 (Bio-Techne) for 3 days, followed by 50 ng/mL rh TGFB1 (Bio-Techne) for a further 3 days in the presence of 100 IU/mL rh IL-2. After cytokine induction, CD8^+^ T cells were purified by magnetic bead isolation according to the manufacturer’s instructions (Miltenyi). Isolated CD8^+^ T-cells were subsequently stained for 20 minutes at room temperature ahead of flow cytometric sorting. Antibodies used: anti-CD45 BV711, anti-CD3 PE-Cy7, anti-CD8 AF700, anti-CD69 PE-Dazzle594 and anti-CD103 FITC. Two dump channels were used: 1 (anti-CD4 and anti-CD56 BUV395) and a second (anti-γ8TCR, anti-Vα7.2 and anti-CD19 BV786) to gate out contaminating cells from the sort strategy. Non-T_RM_-like cells (CD69^-^CD103^-^) and T_RM_-like cells (CD69^+^CD103^+^) were sorted on a FACS Fusion into 50:50 cRPMI:FCS media. Sorted cells were centrifuged at 10,000 rpm for 10 min and resuspend in 350 µL RLT Buffer (Qiagen) and stored at –80 prior to RNA extraction. RNA was extracted using the RNeasy Micro Kit to the manufactures instructions (Qiagen).

RNA quantity and purity were assessed by NanoDrop spectrophotometry (ThermoFisher). Samples with sufficient purity and quantity were submitted to Novogene (Novogene Co., Ltd.) for bulk RNA sequencing.

Polyadenylated mRNA was enriched from total RNA using oligo(dT) magnetic beads. Purified mRNA was fragmented, and first-strand cDNA synthesized using random hexamer priming followed by second-strand cDNA synthesis. After end-repair, A-tailing, and adaptor ligation, libraries were amplified by PCR and purified. Library quality and insert size distribution were verified by Bioanalyzer and quantified by qPCR. Sequencing was performed on the Illumina platform (NovaSeq X Plus Series PE150, paired-end 150 bp). Raw reads underwent quality control, including removal of adaptor sequences, reads containing >10% unknown bases (N), and low-quality reads. Clean reads were aligned to the human reference genome (GRCh38) using a splice-aware aligner. Gene-level read counts were generated using featureCounts, and downstream differential expression analysis was performed using DESeq2. Heatmap analysis of DEGs between non-T_RM_ and induced T_RM_ was performed using ggplot2.

### CPA quantification

Liver collagen content was quantitatively assessed from histological preparations as described previously^27^, using Digital image analysis of the collagen proportionate area (CPA)^28^. In brief, whole slide images were taken of picrosirius red (PSR)-stained slides and areas of fibrosis were identified and segmented from background tissue using a red, green, blue (RGB) threshold. The CPA measurement is the proportionate area of collagen in respect to the area of the whole tissue section, expressed as a percentage.

### Multiparametric flow cytometry

Multiparametric flow cytometry was used for phenotypic and functional analysis. For analysis of the intrahepatic CD8^+^ T cell compartment a strict gating criterion was used throughout. IHL were stained with a blue fixable live/dead dye (Invitrogen; Thermo Fisher Scientific) for the exclusion of dead cells. Doublets, CD45^−^, CD56^+^, CD19^+^, αβTCR^-^ and CD3^+^CD4^+^ cells were also excluded from the analysis (sequential gating strategy; **Supp.Fig.2a**).

Cell surface markers were stained with saturating concentrations of monoclonal antibodies for 30 min at 4 °C in 50% diluted Brilliant Violet buffer (BD Bioscience). To prevent unwanted binding of antibodies to Fc receptors (FcR), an FcR blocking step (Miltenyi Biotech) was included 20% diluted FcR block, 10 min prior to surface antibody labelling. Once surface-stained cells were fixed using Cytofix/Cytoperm (BD Bioscience). Intracellular markers were subsequently stained in a 0.1% saponin-based buffer (Sigma-Aldrich; Merck) containing 0.5% FBS for 30 min at 4 °C. Full details of the monoclonal antibodies, including dilutions and catalogue numbers, are provided in **Supp.Table2**. All samples were acquired in 1x PBS on a 5-laser Cytek Aurora system running SpectraFlow (v.2.2) or on a BD LSR-FortessaX20-SORP system running DIVA (v.8.0.1) and analysed using FlowJo (v.9.9.4 or v.10.8.1; TreeStar; BD Bioscience).

### Preparation of tissues for imaging

4 µm-thick sections were cut from FFPE tissue blocks using a rotary microtome and mounted on to charged glass slides. Prior to staining, tissue sections were dewaxed in xylene and rehydrated in 97% industrial denatured alcohol. Sections were washed in water and microwaved in pre-heated Tris-based or citrate-based antigen retrieval solution (Vector Laboratories) for 30 min.

### Chromogenic immunohistochemistry

Sections were blocked in 2X casein solution (Vector Laboratories) for 10 min. Sections were subsequently incubated overnight with rabbit anti-human HAL (Atlas Antibodies; 1:200) diluted in casein solution at 4°C. Sections were then washed and incubated with a peroxidase-blocking solution (Dako) for 10 min. After blocking sections were washed twice and incubated with ImmPRESS® HRP Universal Antibody (Horse Anti-Mouse/Rabbit IgG) Polymer Detection Kit (Vector Laboratories) for 30 min at R.T. After additional washes, HAL antibody staining was developed using ImmPACT® DAB Substrate Kit, Peroxidase (HRP) according to the manufacturer’s instructions. Sections were then counterstained with Mayers Haematoxylin for 45 seconds at R.T. Haematoxylin staining was developed by briefly washing in hot water, followed by an immediate dehydration in alcohol, and then xylene. Sections were mounted with DPX mounting medium (CellPath) and imaged on a Zeiss Axio Scan.Z1 Slide Scanner controlled by Zen Blue 2012 (Carl Zeiss).

### Immunofluorescence

Sections were blocked in 2X casein or 1% BSA in TBS with 0.1% Tween20 (TBST; Sigma) solution for 10 min. Sections of either decellularised or normal tissue were subsequently incubated overnight with primary antibodies (detailed below) diluted in chosen blocking solution at 4°C. Sections were then washed and incubated for 1 hr with corresponding secondary antibodies (detailed below) diluted in chosen blocking solution at R.T. Sections were washed and treated with TrueView autofluorescence quenching kit (Vector Laboratories) for 5 min, according to the manufacturer’s instructions. Sections were then rinsed in TBST to remove aggregates and mounted with Vectashield Vibrance with DAPI

(Vector Laboratories) for imaged on a Zeiss LSM 880 confocal microscope ZEN Black v2.3 - using the tile scan feature. Sections were imaged using a x40 objective lens using the lowest digital zoom (0.6). 20 tiles were captured per section. Donor tile scans were centred on clear portal triads identified in corresponding serial sections stained with H&E. For cirrhotic ALD sections, images were captured of portal regions, areas of uninvolved non-fibrotic parenchymal tissue and fibrotic scars as defined by serial sections stain with picrosirius red. Decellularised tissue was imaged using the Nikon Eclipse Ti with corresponding NIS-elements AR 5.42.02 software.

The following primary antibodies were used: rabbit anti-human CD103 (Abcam; EPR22590-27; 1 in 500), mouse anti-human E-cadherin (BD Biosciences; 36/E-cadherin; 1 in 100), and mouse anti-human CD8 (Invitrogen; 4B11; 1 in 100). The corresponding secondary antibodies used were: Dylight 488-conjugated horse anti-rabbit IgG (Vector Laboratories), AlexaFluor 546-conjugated goat anti-mouse IgG2a (Invitrogen) and AlexaFluor 647-conjugated goat anti-mouse IgG2b (Invitrogen). All secondaries were diluted 1 in 1000 in 2X casein solution.

### Immunofluorescent cyclic staining via MACSIMA

FFPE-tissue sections were mounted on charged slides in accordance with the MACSwell (Miltenyi Biotech) ‘two imaging frame’ template. Sections were prepared with an equivalent protocol as described above for immunofluorescence imaging until the addition of the blocking solution. From that point, 0.1% triton-x (Sigma Aldrich) was added to 5% goat serum and 1% FBS in PBS for 30 min at R.T. For multiple-plex panels that included unconjugated primary antibodies (detailed in **Supp.Table3**) these were incubated overnight at 4°C in blocking buffer. Sections were then washed with the MACSima running buffer (Miltenyi Biotech for 1 min at RT three times. Cell nuclei were stained 1:50 in DAPI (Miltenyi Biotech) for 10 min at R.T. prior to a final wash cycle in MACSima running buffer. After washing sections were submerged until acquisition.

MACSima is a fully automated instrument that handles both the liquid handling and microscopy. Therefore, fluorescently labelled antibodies were diluted in MACSima running buffer (containing FcR block). Staining volumes were determined by the machine software in a MACSwell deepwell plate (Miltenyi Biotech). ROIs were defined using the DAPI-stain. Staining cycles consisting of the following automated processes: staining, washing, ROI imaging, signal erasure consisting of either photobleaching or removal via REAlease reagents (Miltenyi Biotech) were carried out in accordance with the machine’s protocol.

### Image analysis

Sections were analysed using Zen lite v3.10. The total area of each donor tissue scan was analysed. For cirrhotic ALD sections, areas of uninvolved non-fibrotic parenchymal tissue and fibrotic scars as defined by serial sections stain with picrosirius red and expression of E-cadherin and further confirmed by an independent histopathologist (examples in **Fig.3**; Dr Gary Reynolds, University Hospital Birmingham). These were outlined in Zen lite using the spline contour tool and their area was measured. Portal regions were defined by first identifying the portal triad and drawing a concentric outline 50 µm from this reference. The total number of CD103^+^ T cells, CD8^+^ T cells, and CD103^+^ CD8^+^ were counted/region using the “events” tool in Zen lite. The frequency of each cell type was normalised to area analysed (mm^2^).

CD103 staining intensity was measured on cells with confirmed CD103 and CD8 co-staining. These were outlined using the spline contour tool in Zen lite. Mean intensity values for CD103 staining for each outlined cell was acquired from the “measure” feature in the Zen lite software. To measure the distance of cells from the portal vein, the portal vein was outlined using the spline contour tool. Measurements were then taken using the “line” tool in Zen lite by drawing a line from the centre of the cell nucleus to the nearest edge of the portal vein.

For images captured on the MACSima, these were initially pre-processed and analysed using the machine built-in software MACS iQ view (Miltenyi Biotech). Segmentation was performed based on the nuclei stain. CD8^+^ T-cells (CD45^+^CD3^+^) were identified with a sequential ‘gating strategy’. To quantify the CD103^+^ T_RM_ in a selected ROI, CD8^+^CD103^+^ ‘dual gating’ was used and an automated count performed.

### Isolation of primary hepatic stellate cells (HSCs) and confirmation of αSMA expression

Primary hepatic stellate cells were isolated as previously described according to Rombouts et al., ^57,58^ from healthy, non-fibrotic liver tissues without any clinical indication, or histological evidence of liver disease or fibrotic/cirrhotic resected NAT or explanted liver tissue samples (REC reference 21/WA/0388) by density centrifugation with an Optiprep (Sigma-Aldrich; Merck) gradient. Full details of the HSC donors are included in **Supp.Table5**. Pre-isolated HSC were thawed and cultured in 25 or 75 cm^2^ tissue culture flasks in stellate cell medium (ScienCell Research Laboratories) and maintained at 37 °C, 5% CO_2_, to approximately 80% confluency prior to experimental use. Medium was refreshed twice a week, and cells were passaged (where appropriate) with 0.05% trypsin-EDTA Life Technologies; Thermo Fisher Scientific). Experiments were performed on HSC preparations between passage 3 and 8.

Immunofluorescent imaging of HSCs was performed by seeding 1x10^4^ cells/well in a 96-well blacked-out microplate (Greiner) and cultured at 37 °C, 5% CO_2_ until 70% confluent. Cells were subsequently fixed in 4% PFA for 10 min at R.T., before being washed 3x with TBST. After washing, cells were permeabilised with 0.01% triton-x in TBST for 5 min. HSCs were incubated with mouse anti-αSMA (1:100, Sigma-Aldrich, clone 1A4) for 1 hr at R.T. before being washed 3x and then further incubated with the corresponding AlexaFluor555 goat anti-mouse (Thermo Fisher Scientific) at 3 µg/mL for 1 hr at R.T. Once antibody stained, cell nuclei were stained for imaging with DAPI (1:10,000) for 10 min. HSCs were imaged on the Nikon Eclipse Ti confocal microscope.

### Primary hepatic stellate cell co-culture with PBMC/IHL

For PBMC co-culture experiments, freshly isolated PBMC from healthy donors at 3 x 10^5^ cells/well were treated with either 50 ng/mL recombinant human (rh) IL-15 or 0.5 µg/mL immobilised anti-CD3 for 3 d in complete RPMI-1640 (cRPMI; containing 10% FBS, 100 IU/mL penicillin-streptomycin, 20 mM HEPES, 0.5 mM sodium pyruvate, MEM non-essential amino acids, MEM essential amino acids and 50 µM β−mercaptoethanol; all Life Technologies; Thermo Fisher Scientific) containing 100 IU/mL recombinant human IL-2 (PeproTech) before exposure to HSCs. During the IL-15 or TCR-activation period, on d 2, pHSC were detached using trypsin-EDTA and reseeded at a density of 0.2 x 10^5^ cells per well in stellate cell medium in a 24-well plate and left to adhere. On d 3 PBMC were washed in cRPMI and transferred for co-culture with pre-seeded HSCs for a further 3 days at 37 °C. Where indicated the addition of 1 µg/mL anti-TGFβ (BioTechne) was added to the co-cultures.

For the IHL co-culture experiments, HSC were detached using trypsin-EDTA and reseeded at a density of 0.2 x 10^5^ cells per well in stellate cell medium in a 24-well plate and left to adhere. Subsequently HSC were co-cultured with the addition of 1 x 10^6^ freshly isolated or thawed IHL in complete RPMI-1640 containing 100 IU/mL recombinant human IL-2 for 16 hr, at 37 °C. Where indicated HSCs were either preincubated with 5 µg/mL anti-PD-L1/-L2 antibodies (Thermo Fischer Scientific) for 1 hr (after incubation unbound antibody was washed off using 1x PBS) was added during co-culture.

### Preparation and use of 3D decellularised liver scaffolds

For the preparation of decellularised scaffolds, liver tissue was obtained through the TAPb Initiative as above. Acellular 3D biological scaffolds “cubes” were generate using previously published protocols^31,32^. Briefly, frozen 0.5mm^3^ liver cubes were thawed, agitated with cycles of deionized water (Merck-Millipore) containing 3% sodium deoxycholate (Sigma-Aldrich; Merck), 0.5% sodium dodecyl sulphate (Sigma-Aldrich; Merck), 0.3% Triton X-100 (Thermo Fisher Scientific) and 4.5% sodium chloride (Sigma-Aldrich; Merck) and 1x PBS until the tissue was translucent from the dissolution of cells. The absence of cells in the ECM protein scaffolds was confirmed by the absence of DNA material quantified using the DNeasy Blood and Tissue Kit (Qiagen) and H&E stains. Scaffolds generated with this protocol have previously been confirmed by proteomics and immunohistochemical analysis to express natural ECM including collagen-I, -III and -IV, fibronectin, laminin, lumican, mimecan and vitronectin^30^. Prior to use 3D decellularised human liver scaffolds were sterilized using 0.1% peracetic acid (Sigma-Aldrich, St. Louis, USA) and left overnight in cRPMI containing 100 IU/mL recombinant IL-2.

For co-culture experiments 3x 3D acellular ECM scaffolds (from the same tissue donor) were repopulated with freshly isolated PBMC (1 x 10^6^) in cRPMI in the presence 100 IU/mL recombinant human IL-2 for 3 d in 24-well round-bottom plates. After culture, the 3D scaffolds (and cells cultured in the absence of a 3D scaffold) were digested for 30 min with HBSS^+/+^ containing 0.002% DNase I and 0.02% collagenase IV to recover cells for phenotypic analysis as described in the ‘Multiparametric flow cytometry’ section.

### Assessment of T_RM_ phenotype and function

To assess the functionality of intrahepatic CD8^+^ T cells *ex vivo*, 1 x 10^6^ IHL were stimulated with 1 µg/mL immobilised anti-CD3 and 5 µg/mL soluble anti-CD28 for 4 hr at 37 °C in the presence of 1 µg/mL brefeldin A (Sigma-Aldrich; Merck) and 0.66 µL/mL BD Golgi Stop containing monensin (BD Bioscience). Functionality was assessed by intracellular cytokine production as described in the ‘Multiparametric flow cytometry’ section.

To assess the functionality of hepatic CD8^+^ T cells after co-culture with HSC (described in the ‘*Isolation of primary human hepatic stellate cells and co-culture with IHL’* sections above) Isolated IHL were stimulated with 1x10^6^/mL Human T-Activator CD3/CD28 Dynabeads (Thermo Fisher Scientific) in cRPMI supplemented with 100 IU/mL IL-2, 1 µg/mL brefeldin A and 0.66 µL/mL Golgi Stop for 4 hr or overnight with or without HSC. Functionality was assessed by intracellular cytokine production as described in the ‘Multiparametric flow cytometry’ section.

### Statistical analyses

Statistical analyses were performed in Excel (v. 16.89.1) and Prism (GraphPad v.10.3.1) using the appropriate tests (Mann–Whitney *t-*tests, Wilcoxon signed-rank *t-*tests, Kruskal–Wallis tests (ANOVA) with Dunn’s post hoc test for multiple comparisons between each group) as indicated in the legends. These tests were performed as two-tailed tests, with significant differences marked on all figures. Error bars represent standard error of the mean (S.E.M.). Significance levels are marked on all figures. Relevant visualisations were created using Python (v.3.6) and the seaborn statistical visualisation package.

## Acknowledgments

We are extremely grateful for, and would like to take this opportunity to thank, all the patients and control volunteers who participated in this study. We would also like to thank the clinical staff who helped with study participant recruitment, sample acquisition and provision – particularly members of the Tissue Access for Patient Benefit Initiative (TAPB) at The Royal Free Hospital. We would also like to thank the staff in the Flow Cytometry Core Facility (Division of Infection & Immunity; IIT).

## Author contributions

LJP conceived the study and obtained funding, GEF, BHJ, SPD, PR, LJP designed experiments, GEF, DRB, SK, BHJ, KR, GP, AM, AH, SPD carried out experiments, GEF, DRB, BHJ, SK, KR, GP analysed data, SPD, GL, WA, AH, KB, KR, AQ, MKM provided clinical samples, experimental protocols and/or valuable expertise, GEF, LJP wrote the manuscript, all other authors provided critical review of the manuscript.

## Data and materials availability

[to complete after initial submission]

---

**Supplementary Figure 1.** *Identification of CD103^+^ T_RM_ and assessment of frequencies in relation to clinical parameters.* **a**) Representative flow cytometry gating strategy to define hepatic CD8^+^ T-cells with sequential exclusion of; debris by forward (FSC-A) and side scatter (SSC-A), doublets, dead cells, CD45^-^, CD3^-^, CD56^+^, αβTCR^-^, CD3^+^CD4^+^. **b)** Frequency of CD69^+^CD103^+^ T_RM_ as a proportion of total live intrahepatic leukocytes (IHL; defined as CD45^+^) in healthy, non-CLD livers (Controls; samples without clinical indication, or histological evidence of liver disease [n=89]) and CLD livers (n=56). **c**) Correlation of the frequency of hepatic CD103^+^ T_RM_ with age (years). **d**) Frequency of CD103^+^ T_RM_ identified in the liver in each cohort - healthy (no evidence of liver disease) or CLD - categorised by sex (where known). Correlation of the frequency of hepatic CD103^+^ T_RM_ with serum **e)** alanine transaminase (ALT) and **f)** aspartate transferase (AST). **g)** Frequency of CD103^+^ T_RM_ (top) and CD103^-^T_RM_ (bottom) categorised by independent histological assessment using the ISHAK scoring system. **h)** Representative images showing serial a section of a non-fibrotic, control liver section stained with haematoxylin and eosin (H&E). **i**) Summary data depicting the mean florescence (arbitrary units; AU) of E-cadherin in the fibrotic scar compared to the uninvolved, non-fibrotic parenchymal region (uninvolved) as determined by confocal microscopy of cirrhotic ALD tissue sections (n=7). Violin plots show median ± quartiles. Each tissue sample (denoted as individual dots) was stained and processed independently. Error bars represent mean ± S.E.M.

**Supplementary Figure 2.** *Confirmation of HSC αSMA expression, CD103 expression on activated CD8^+^ T cells after co-culture and RNA-atlas analyses*. **a**) Representative images showing alpha smooth muscle (αSMA; red) expression by primary hepatic stellate cells (HSCs) isolated from a cirrhotic tissue sample by immunofluorescence. **b**) Representative flow cytometric plots showing cirrhotic HSC expression of intracellular LAP-TGFβ compared to a fluorescence minus control (FMO; top) and summary data depicting the mean fluorescence intensity of LAP-TGFβ staining across all batches of control, non-fibrotic HSCs (n=7) and cirrhotic HSCs (n=8). **c**) Summary data depicting percentage CD103 expression on peripheral IL-15-activated CD8^+^ T-cells in media alone (white; n=12), after recombinant TGFβ1 exposure (grey; n=12) or co-culture with primary HSCs (purple; n=12). **d)** Summary data depicting expression of CD11a, CD49b and CD29 on CD103^+^ T_RM_ identified in tissue samples without any clinical indication, or histological evidence of liver disease (Control: blue) compared to a combined cohort of samples with mild-moderate fibrosis and advanced fibrosis (Fibrotic; green). **e**) UMAP visulisation of scRNAseq data (107,488 hepatic CD8^+^ T-cells; 77 patients) identifying eight clusters, coloured and labelled by marker gene expression profiles. **f+g**) Dotplots depicting the mean expression of highly expressed marker genes and the percentage of T-cells expressing them to define each cluster identified in **e**. Error bars show mean ± S.E.M. Violin plots show median ± quartiles. Each tissue sample was stained and processed independently.

**Supplementary Figure 3.** *CD103^+^ T_RM_ staining and analysis of a range of co-inhibitory and co-stimulatory markers.* Representative flow cytometry plots for **a**) PD1 and **b**) CD28 and LAG3 on CD69^-^CD103^-^ liver-infiltrating and CD103^+^ T_RM_ identified in a tissue sample without any clinical indication, or histological evidence of liver disease (top; healthy, non-fibrotic) compared to a sample with fibrosis (bottom; CLD). **c**) tSNE plot visualising expression of a range of activatory, co-inhibitory and co-stimulatory receptors expressed by healthy (n=9) and CLD (n=10) CD103^+^ T_RM_ – individual feature plots depict the expression profile of each marker individually. **d**) Percentage of phenotypic signatures CD103^+^ T_RM_ evaluated by conventional flow cytometry gating in samples with or without evidence of fibrosis showing PD-1^hi^LAG3^hi^TIM3^hi^ and TIM3^hi^CD28^-^. Each IHL sample was processed and stained independently. Violin plots show median ± quartiles.

**Supplementary Figure 4.** *HSC flow cytometry gating controls.* **a**) Representative flow cytometry plots depicting expression of PD-L1 on HSCs isolated from a non-fibrotic, control or cirrhotic liver compared to a fluorescence minus one control (FMO; top). **b**) Representative flow cytometry plots depicting expression of PD-L1 on the surface of HSC post incubation for 1h with either PD-L1/L2 neutralising antibodies or an isotype control (IgGκ1).

