## Supplementary Files for "Human hepatic stellate cells orchestrate the accumulation and function of CD103^+^ tissue-resident CD8^+^ T-cells in liver fibrosis"

### Supplementary Figure 1

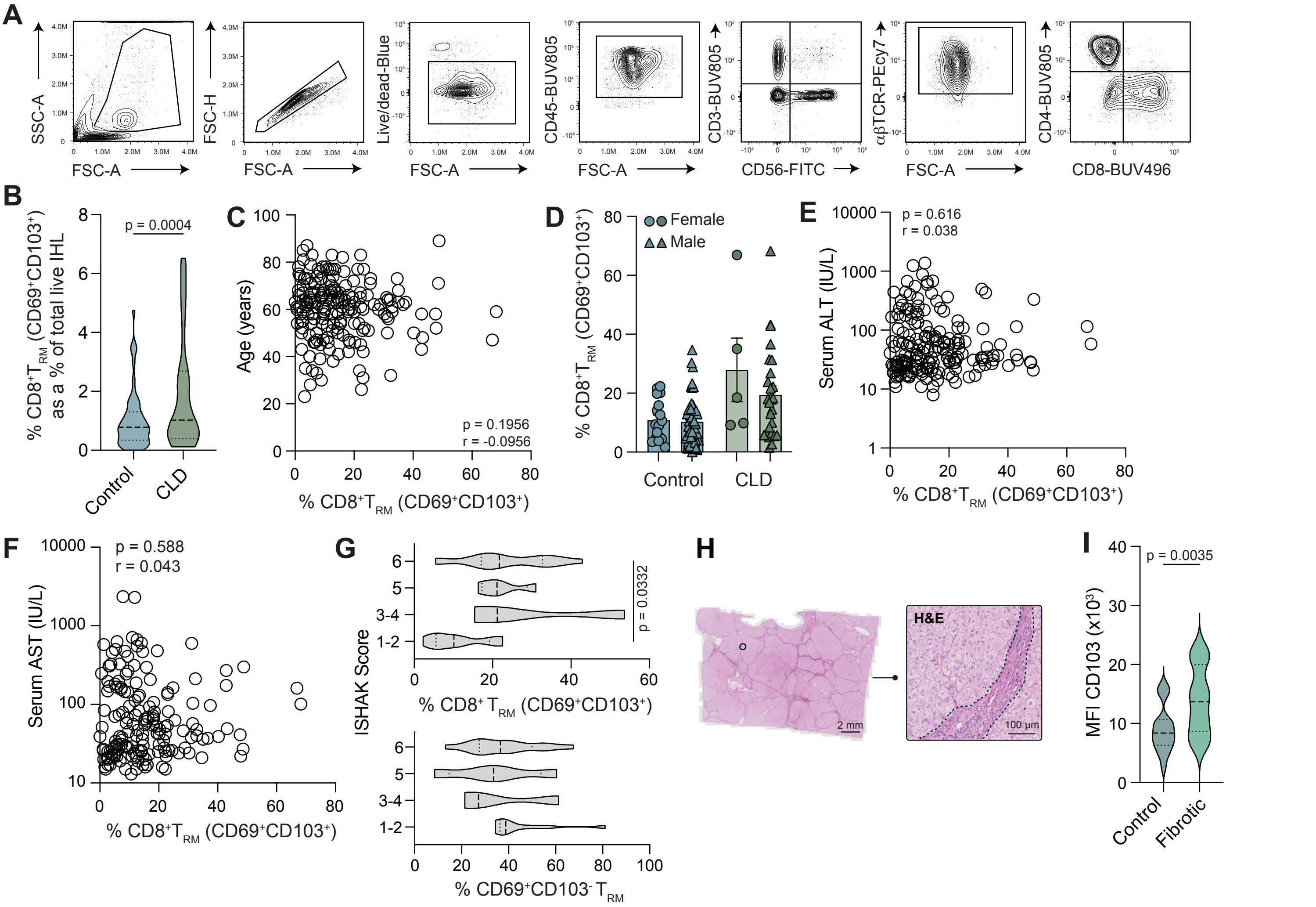


**Supplementary Figure 1**: *Identification of CD103$^+$ $T_{RM}$ and assessment of frequencies in relation to clinical parameters.* **a**) Representative flow cytometry gating strategy to define hepatic CD8$^+$ T-cells with sequential exclusion of; debris by forward (FSC-A) and side scatter (SSC-A), doublets, dead cells, CD45$^-$, CD3$^-$, CD56$^+$, αβTCR$^-$, CD3$^+$CD4$^+$. **b**) Frequency of CD69$^+$CD103$^+$ $T_{RM}$ as a proportion of total live intrahepatic leukocytes (IHL; defined as CD45$^+$) in healthy, non-CLD livers (Controls; samples without clinical indication, or histological evidence of liver disease [n=89]) and CLD livers (n=56). **c**) Correlation of the frequency of hepatic CD103$^+$ $T_{RM}$ with age (years). **d**) Frequency of CD103$^+$ $T_{RM}$ identified in the liver in each cohort - healthy (no evidence of liver disease) or CLD - categorised by sex (where known). Correlation of the frequency of hepatic CD103$^+$ $T_{RM}$ with serum **e**) alanine transaminase (ALT) and **f**) aspartate transferase (AST). **g**) Frequency of CD103$^+$ $T_{RM}$ (top) and CD103$^-$ $T_{RM}$ (bottom) categorised by independent histological assessment using the ISHAK scoring system. **h**) Representative images showing serial a section of a non-fibrotic, control liver section stained with haematoxylin and eosin (H&E). **i**) Summary data depicting the mean florescence (arbitrary units; AU) of E-cadherin in the fibrotic scar compared to the uninvolved, non-fibrotic parenchymal region (uninvolved) as determined by confocal microscopy of cirrhotic ALD tissue sections (n=7).

Violin plots show median ± quartiles.
Each tissue sample (denoted as individual dots) was stained and processed independently.
Error bars represent mean ± S.E.M.

### Supplementary Figure 2

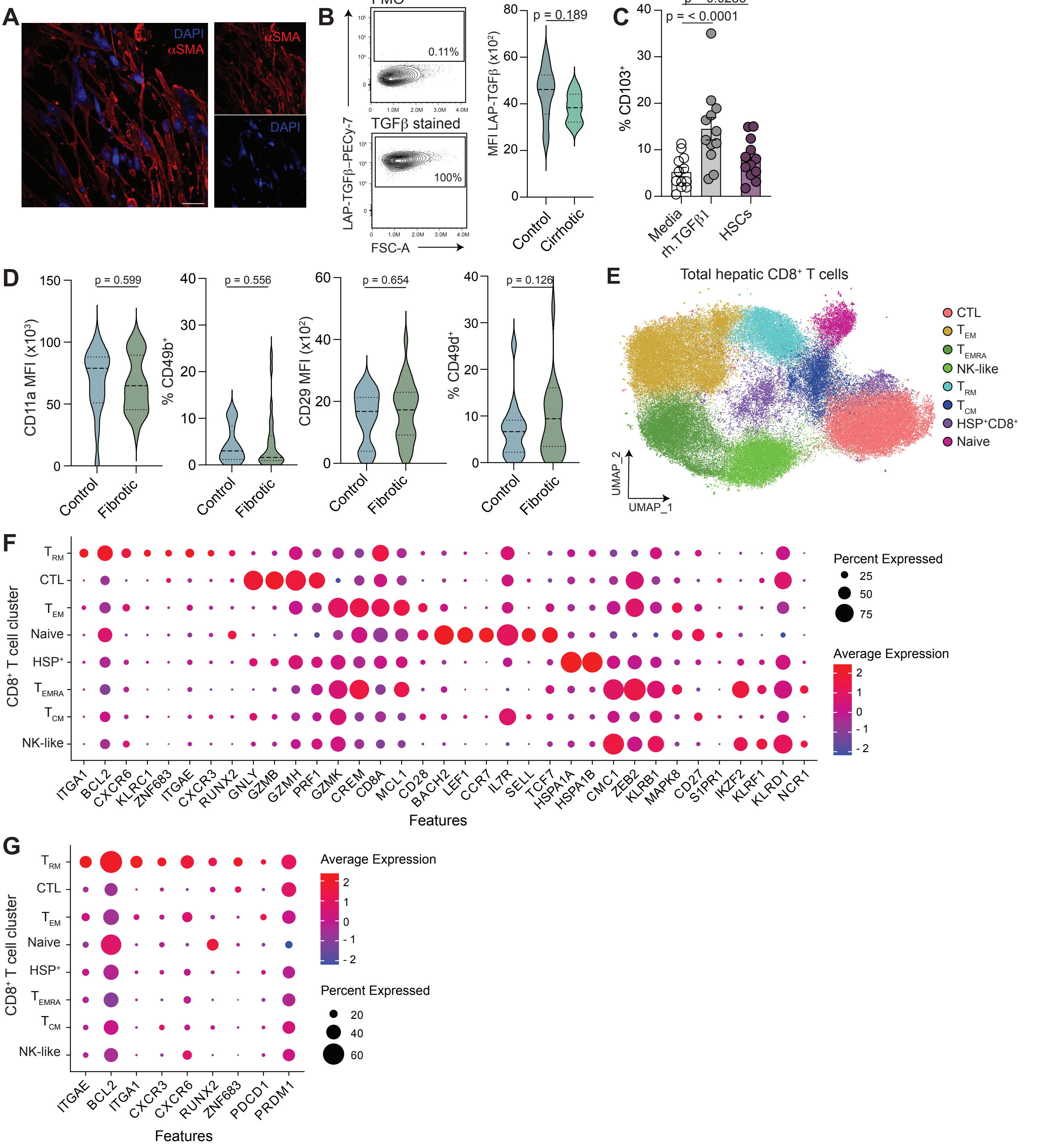


**Supplementary Figure 2:** *Confirmation of HSC αSMA expression, CD103 expression on CD8⁺ T cells after co-culture and RNA-atlas analyses.* **a**) Representative images showing alpha smooth muscle (αSMA; red) expression by HSCs isolated from a cirrhotic tissue sample by immunofluorescence. **b**) Representative flow cytometric plots showing cirrhotic HSC expression of intracellular LAP-TGFβ compared to a fluorescence minus control (FMO; top) and summary data depicting the mean fluorescence intensity of LAP-TGFβ staining across all batches of control, non-fibrotic HSCs (n=7) and cirrhotic HSCs (n=8). **c**) Summary data depicting percentage CD103 expression on peripheral IL-15-activated $CD8^+$ T-cells in media alone (white; n=12), after recombinant TGFβ1 exposure (n=12; grey) or co-culture with HSCs (n=12; purple). **d**) Summary data depicting expression of CD11a, CD49b and CD29 on $CD103^+$ $T_{RM}$ identified in tissue samples without any clinical indication, or histological evidence of liver disease (Control: blue) compared to a combined cohort of samples with mild-moderate fibrosis and advanced fibrosis (Fibrotic; green). **e**) UMAP visulisation of scRNAseq data (107,488 hepatic $CD8^+$ T-cells; 77 patients) identifying 8 clusters, coloured and labelled by marker gene expression profiles. **f+g**) Dotplots depicting the mean expression of highly expressed marker genes and the percentage of T-cells expressing them to define each cluster identified in **e**.
Error bars show mean ± S.E.M.
Violin plots show median ± quartiles.

### Supplementary Figure 3

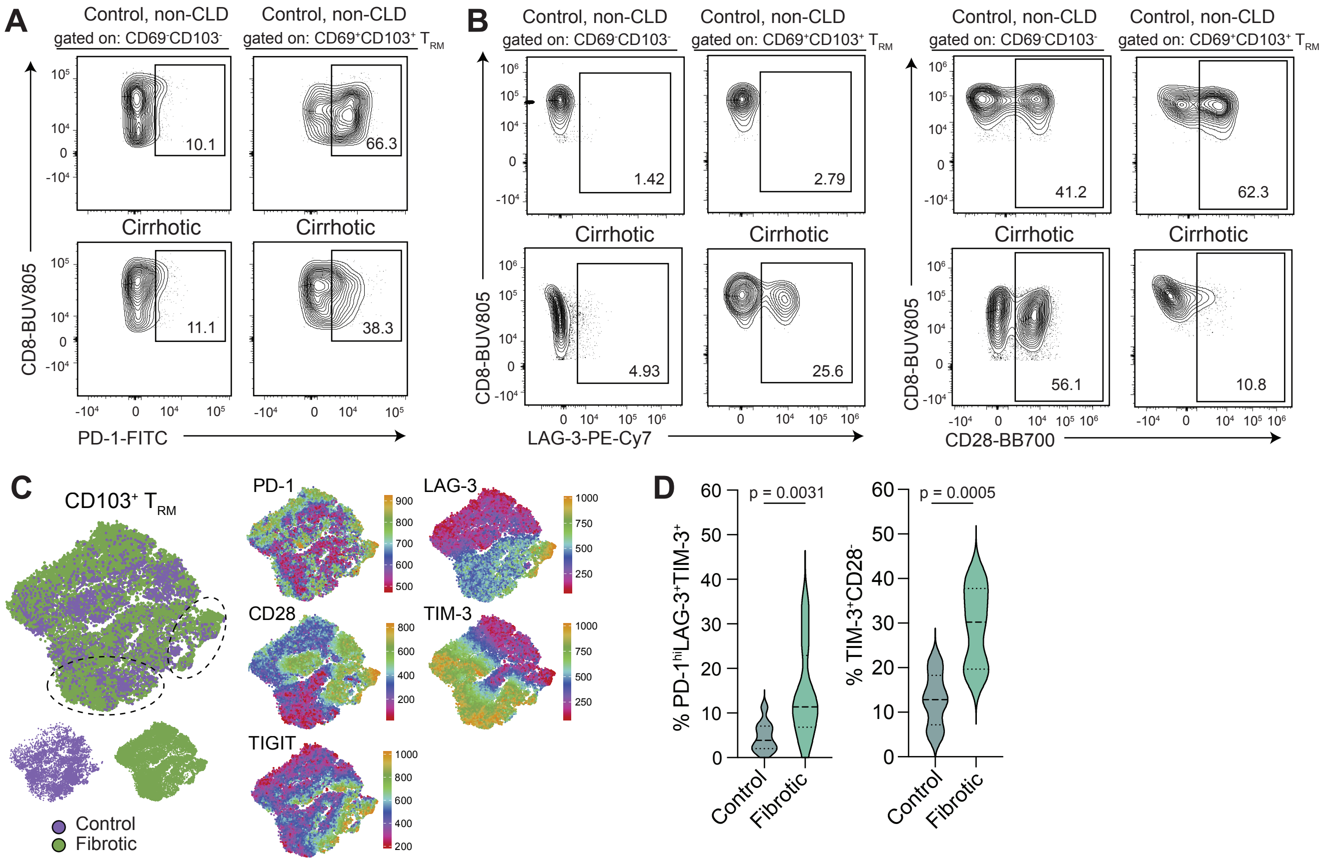


**Supplementary Figure 3:** *CD103⁺ $T_{RM}$ staining and analysis of a range of co-inhibitory and co-stimulatory markers.* Representative flow cytometry plots for **a**) PD1 and **b**) CD28 and LAG3 on CD69⁻CD103⁻ liver-infiltrating and CD103⁺ $T_{RM}$ identified in a tissue sample without any clinical indication, or histological evidence of liver disease (top; healthy, non-fibrotic) compared to a sample with fibrosis (bottom; CLD). [c]) tSNE plot visualising expression of a range of activatory, co-inhibitory and co-stimulatory receptors expressed by healthy (n=9) and CLD (n=10) CD103⁺ $T_{RM}$ – individual feature plots depict the expression profile of each marker individually. **d**) Percentage of phenotypic signatures CD103⁺ $T^{RM}$ evaluated by conventional flow cytometry gating in samples with or without evidence of fibrosis showing PD-$1^{hi}$LAG$3^{hi}$TIM$3^{hi}$ and TIM$3^{hi}$CD$28^{-}$.
Each IHL sample was processed and stained independently.
Violin plots show median ± quartiles.

### Supplementary Figure 4

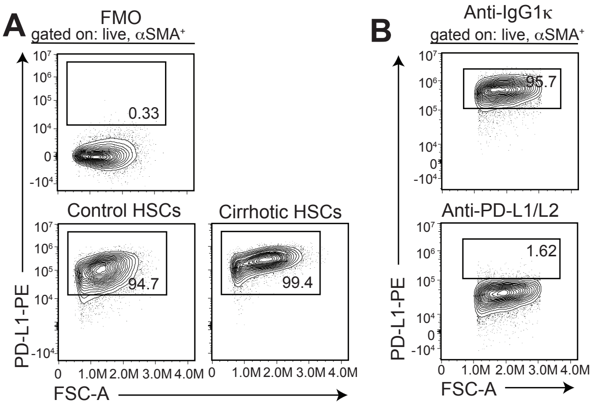


**Supplementary Figure 4:** *HSC flow cytometry gating controls.* **a**) Representative flow cytometry plots depicting expression of PD-L1 on HSCs isolated from a non-fibrotic, control or cirrhotic liver compared to a fluorescence minus one control (FMO; top). **b**) Representative flow cytometry plots depicting expression of PD-L1 on the surface of HSC post incubation for 1h with either PD-L1/L2 neutralising antibodies or an isotype control (IgGκ1), confirming blocking.

**Supplementary Table 1:** Clinical paramters defining the intrahepatic leukocyte cohort

| | | | CLD | | HCC | |
|---|---|---|---|---|---|---|
| | | CRCLM (n=89) | met-ALD (n=26) | MASH/MASLD (n=22) | met-ALD (n=7) | MASH/MASLD (n=14) |
| **Age** | | | | | | |
| **(yrs)** | median | 63 | 59 | 64 | 60 | 67 |
| | ICR | 25 (49-71) | 12 (51-63) | 10 (59-69) | 17 (46-63) | 10 (61-71) |
| **Sex** | | | | | | |
| **Female** | n (%) | 24 (27%) | 3 (12%) | 0 (0%) | 0 (0%) | 0 (0%) |
| **Male** | n (%) | 65 (73%) | 23 (88%) | 22 (100%) | 7 (100%) | 14 (100%) |
| **Clinical parameters** | | | | | | |
| **ALT (IU/L)** | median (+range) | 46 (8-1373) | 35 (16-63) | 46.5 (13-202) | 35 (25-57) | 44 (16-202) |
| **AST (IU/L)** | median (+range) | 40 (15-2304) | 49 (24-111) | 53 (15-230) | 44 (24-97) | 39 (16-267) |
| **Bilirubin (µmol/L)** | median (+range) | 11 (2-154) | 32 (4-652) | 10 (4-72) | 25 (6-42) | 8.5 (4-20) |
| **Platelet count** | median (+range) | 237.5 (22-436 | 90.5 (46-334) | 201.5 (50-370) | 95 (61-334) | 224 (92-370) |
| **Lymphocyte count** | median (+range) | 1.32 (0.34-3.32) | 1.2 (0.54-2.45) | 1.38 (0.65-3) | 0.97 (0.54-1.39) | 1.57 (0.95-3) |
| **FIB-4 score** | median (+range) | 1.90 (0.39-15.6) | 4.44 (0.44-11.4) | 2.63 (0.93-15.2) | 4.57 (0.44-7.1) | 1.59 (0.93-8.8) |
| **Fibrosis assessment** | | | | | | |
| **Healthy, non-fibroti** | n (%) | 69 (78%) | 0 (0%) | 1 (4%) | 0 (0%) | 0 (0%) |
| **Fibrotic** | n (%) | 20 (22%) | 0 (0%) | 10 (46%) | 1 (14%) | 8 (67%) |
| **Cirrhotic** | n (%) | 0 (0%) | 26 (100%) | 11 (50%) | 6 (86%) | 6 (27%) |

**Supplementary Table 2:** Details of antibodies used for multiparametric flow cytometery

| Marker | Fluorphore | Clone | Company | Dilution |
|---|---|---|---|---|
| CD3 | BUV395 | UCHT1 | BD Biosciences | 0.25/100 |
| CD8 | BUV496 | RPA-T8 | BD Biosciences | 0.5/100 |
| CD49b | BUV563 | AK-7 | BD Biosciences | 1/100 |
| CD49e | BUV615 | IIA1 | BD Biosciences | 1/100 |
| CD56 | BUV737 | NCAM16.2 | BD Biosciences | 0.25/100 |
| CD45 | BUV805 | HI30 | BD Biosciences | 0.25/100 |
| CD29 | SB436 | TS2/16 | ThermoFischer | 1/100 |
| CD44 | V450 | G44-26 | BD Biosciences | 1/100 |
| HLA-DR | BV570 | L243 | Biolegend | 1/100 |
| CD38 | BV650 | HB-7 | Biolegend | 0.5/100 |
| TIGIT | BV711 | TgMab-2 | BD Biosciences | 1/100 |
| PD-1 | BV750 | EH12.2H7 | Biolegend | 0.5/100 |
| CD19 | BV786 | HIB19 | BD Biosciences | 0.25/100 |
| CD49d | BB515 | 9F10 | BD Biosciences | 1/100 |
| CD4 | Spark Blue 550 | SK3 | Biolegend | 0.25/100 |
| CD49a | PE | REA1106 | Miltenyi | 0.5/100 |
| CD69 | PE-Dazzle594 | FN50 | Biolegend | 0.5/100 |
| ab-TCR | PECy5 | IP26 | Biolegend | 0.25/100 |
| CD11a (aL) | PECy7 | HI111 | Biolegend | 1/100 |
| CXCR6 | AF647 | K041E5 | Biolegend | 1/100 |
| CD103 | APC-Cy7 | BER-ACT8 | Biolegend | 1/100 |
| CD28 | BB700 | L293 | BD Biosciences | 1/100 |
| Tim3 | BB515 | 7D3 | BD Biosciences | 1/100 |
| LAG3 | PE-Cy7 | 3DS223H | Invitrogen | 1/100 |
| TGFbRII | FITC | W17055E | Biotechne | 1/100 |
| IFNg | V450 | B27 | BD Biosciences | 2/100 |
| IFNg | APC | 4S.B3 | Biolegend | 2/100 |
| TNFa | FITC | MAb11 | Biolegend | 2/100 |
| IL-2 | BB700 | MQ1-17H12 | BD Biosciences | 2/100 |
| IL-2 | BV711 | 5344 111 | BD Biosciences | 2/100 |
| CD107a | PE | H4A3 | BD Biosciences | 2/100 |
| CD107a | BV605 | H4A3 | Biolegend | 2/100 |
| Granzyme B | AF700 | GB11 | BD Biosciences | 1/100 |
| PD-L1 | PE | 29E.2A3 | Biolegend | 1/100 |
| aSMA | eFlour 660 | 1A4 | Invitrogen | 1/100 |
| TGFb1 | PE-Cy7 | S200006A | Biolegend | 1/100 |
| LIVE/Dead Blue | N/A | N/A | Invitrogen | 1/1000 |
| LIVE/Dead Near IR | N/A | N/A | Invitrogen | 1/1000 |

**Supplementary Table 3:** Primary hepatic stellate cell (HSCs) donor details where avaliable

| batch_id | source | sex | serum_alt | serum_bilirubin | serum_ast | platelet_count | wc_count | lymphs_count | fibrosis_assessment |
|---|---|---|---|---|---|---|---|---|---|
| 030114 | HCC | | | | | | | | cirrhotic |
| 040314 | HCC | | | | | | | | cirrhotic |
| 140113 | CRCLM | | | | | | | | healthy |
| 160513 | unknown | | | | | | | | unknown |
| 160516 | CCA | F | 200 | 84 | 164 | 201 | 7 64 | 3 28 | healthy |
| 010616 | CHB (explanted) | | | | | | | | cirrhotic |
| 200716 | CRCLM | M | 29 | 9 | 29 | 323 | 7 | 2 46 | healthy |
| 141119 | CRCLM | F | 27 | 7 | 28 | 444 | 6 86 | 2 03 | healthy |
| 140521 | CCA | F | 29 | 4 | 33 | 253 | 8 98 | 2 89 | unknown |
| 040621 | CRCLM | M | 27 | 5 | 33 | 230 | 9 4 | 1 48 | healthy |
| 110621 | HCC | M | 9 | 9 | 45 | 79 | 4 89 | 0 87 | cirrhotic |
| 220621 | CRCLM | | | | | | | | unknown |
| 030223 | ALD | F | 151 | 31 | | 172 | 8 69 | 0 98 | cirrhotic |
| 201023 | ALD | M | 17 | 4 | | 274 | 4 31 | 0 66 | cirrhotic |
| 181024 | MASH | M | 13 | 6 | 15 | 164 | 7 47 | 0 75 | cirrhotic |
| 211124 | ALD | M | 16 | 11 | 24 | 276 | 10 43 | 2 45 | cirrhotic |
| 281015 | CRCLM | | | | | | | | unknown |
| hscA | HCC | F | | | | | | | fibrosis (F2) |
| hscB | CCA | M | | | | | | | cirrhotic (F4) |
| hscC | unsuitable for transplantation | | | | | | | | healthy (F0) |
| hscD | unknown (purchased from Lonza) | | | | | | | | unknown |
| hscE | unknown (purchased from Lonza) | | | | | | | | unknown |
| hscF | HCC | F | | | | | | | healthy (F0) |
| 270616 | CRCLM | | | | | | | | unknown |

**Supplementary Figure 4:** Details of MACSIMA imaging antibodies

| Marker | Supplier | Clone | Flourochrome |
|---|---|---|---|
| CD3 | Miltenyi | REA1151 | |
| CD8a | Miltenyi | REA1024 | FITC |
| CD326 (EpCAM) | Miltenyi | REA1311 | PE |
| CD324 (E-Cadherin) | Miltenyi | REA1319 | PE |
| CD45 | Biolegen | H130 | AF488 |
| CD103 | Abcam | EPR4166(2) | AF647 |
| αSMA | Sigma-Aldrich | 1A4 | unconjugated |
| TGFβ | ThermoFisher | TB21 | unconjugated |
| collagen-I | Miltenyi | REAL958 | APC |
| FAP | R&D | 427819 | APC |
| collagen-IV | Dako | CIV22 | unconjugated |
